# Cell fate specification modes shape transcriptome evolution in the highly conserved spiral cleavage

**DOI:** 10.1101/2024.12.25.630330

**Authors:** Yan Liang, Jingcheng Wei, Yue Kang, Allan M. Carrillo-Baltodano, José M. Martín-Durán

**Author notes:** Correspondence: José M. Martín-Durán.

## Abstract

Early animal development can be remarkably variable, influenced by lineage-specific reproductive strategies and adaptations. Yet, early embryogenesis is also strikingly conserved in some groups, like Spiralia (e.g., snails, clams, and many marine worms). In this large clade, a shared cleavage program –– the so-called spiral cleavage –– and similar cell lineages are ancestral to at least seven animal phyla. Why early development is so conserved in specific groups and plastic in others is not fully understood. Here, we investigated two annelid species –– *Owenia fusiformi*s and *Capitella teleta* –– with spiral cleavage but different modes of specifying their primary progenitor cells. By generating high-resolution transcriptomic time courses from the oocyte and fertilisation until gastrulation, we show transcriptional dynamics are markedly different between these species during spiral cleavage and instead reflect their distinct timings of embryonic organiser specification. However, the end of cleavage and gastrulation exhibit high transcriptomic similarity. At these stages, orthologous transcription factors share gene expression domains, suggesting this period is a previously overlooked mid-developmental transition in annelid embryogenesis. Together, our data reveal hidden developmental plasticity in the genetics underpinning spiral cleavage, indicating an evolutionary decoupling of morphological and transcriptomic conservation during early animal embryogenesis.

## Introduction

Upon fertilisation, animal zygotes enter a phase of intense cell division called cleavage, resulting in an embryo –– a blastula –– made of developmentally committed cells that segregate into the main germ layers during gastrulation (1, 2). As a foundational stage for subsequent development, broadly conserved cleavage modes are recognised in animals (1, 2). In the holoblastic radial cleavage, the zygote and its daughter cells divide completely and maintain a radially symmetrical arrangement, at least during the early rounds of cell division. This cleavage mode is widespread, observed throughout animal phylogeny, and likely ancestral (3). However, specific clades, such as tunicate chordates and acoel flatworms, have evolved idiosyncratic and highly stereotypic cleavage programmes, like bilateral cleavage and duet spiral cleavage, respectively (4, 5). Indeed, the initial steps of animal embryogenesis can vary highly, often influenced by the reproductive strategies of each animal lineage. This is most obvious in animals whose eggs have large amounts of yolk, such as fishes and birds, insects, and cephalopods (1, 2), where complete cytokinesis during cell division does not occur. Deviations in early cleavage can also be more subtle. For example, although starfish and sea urchins share holoblastic radial cleavage, only the latter form a set of vegetal micromeres that contribute to forming the larval skeleton, a unique trait of this echinoderm lineage (6). How do distinct cleavage modes evolve? How do novelties appear during early embryogenesis, and by what genetic and developmental mechanisms?

Spiral cleavage is an ancient and highly conserved early developmental program found in at least seven major animal groups within Spiralia (or Lophotrochozoa, by some authors), one of the three largest branches of bilaterally symmetrical animals (7–9). The alternating shift of the mitotic spindle along the animal-vegetal embryonic axis from the 8-cell stage onwards characterises this cleavage mode and results in a spiral-like arrangement of the animal-pole blastomeres when viewed from above, hence the name of this early embryogenesis (10–12). In addition to the conserved pattern of cell divisions, embryos with spiral cleavage also show broadly conserved cell lineages (13, 14). Thereby, equivalent blastomeres in closely and more distantly phylogenetically related species often act as progenitors of similar cell types, tissues, and organs. Despite the conservation of cleavage patterns and cell lineages, embryos with spiral cleavage exhibit two markedly different strategies to specify the primary cell lineages and establish their axial patterning (Figure 1) (10–12, 15). In the so-called equal (or conditional) spiral cleavage, bilateral symmetry is established with the inductive specification of a blastomere –– the so-called 4d micromere –– that acts as an embryonic organiser at the 5^th^ or 6^th^ round of cell division (the 32-or 64-cell stages, depending on the species) (Figure 1A–C) (15). In molluscs and annelids, the FGF receptor pathway and ERK1/2 transducing cascade controls this process (10, 16–18). However, symmetry breaking occurs much earlier in many molluscan and annelid embryos. In these species, spiral cleavage is regarded as unequal (or autonomous) because the asymmetric segregation of maternal determinants into a larger cell by the 2^nd^ round of cell division (the 4-cell stage) defines the posterodorsal fate and the progenitor lineage of the embryonic organiser (Figure 1D–F) (19, 20). While equal spiral cleavage occurs in all major animal clades with spiral cleavage and is considered the ancestral condition, the unequal mode has evolved independently multiple times (19). Therefore, spiral cleavage, with overall conserved cell division patterns and cell lineages but recursively evolved distinct modes of cell fate specification and axial patterning, is an ideal system to explore the developmental and evolutionary mechanisms that generate phenotypic change during early animal embryogenesis.

**Figure 1.**
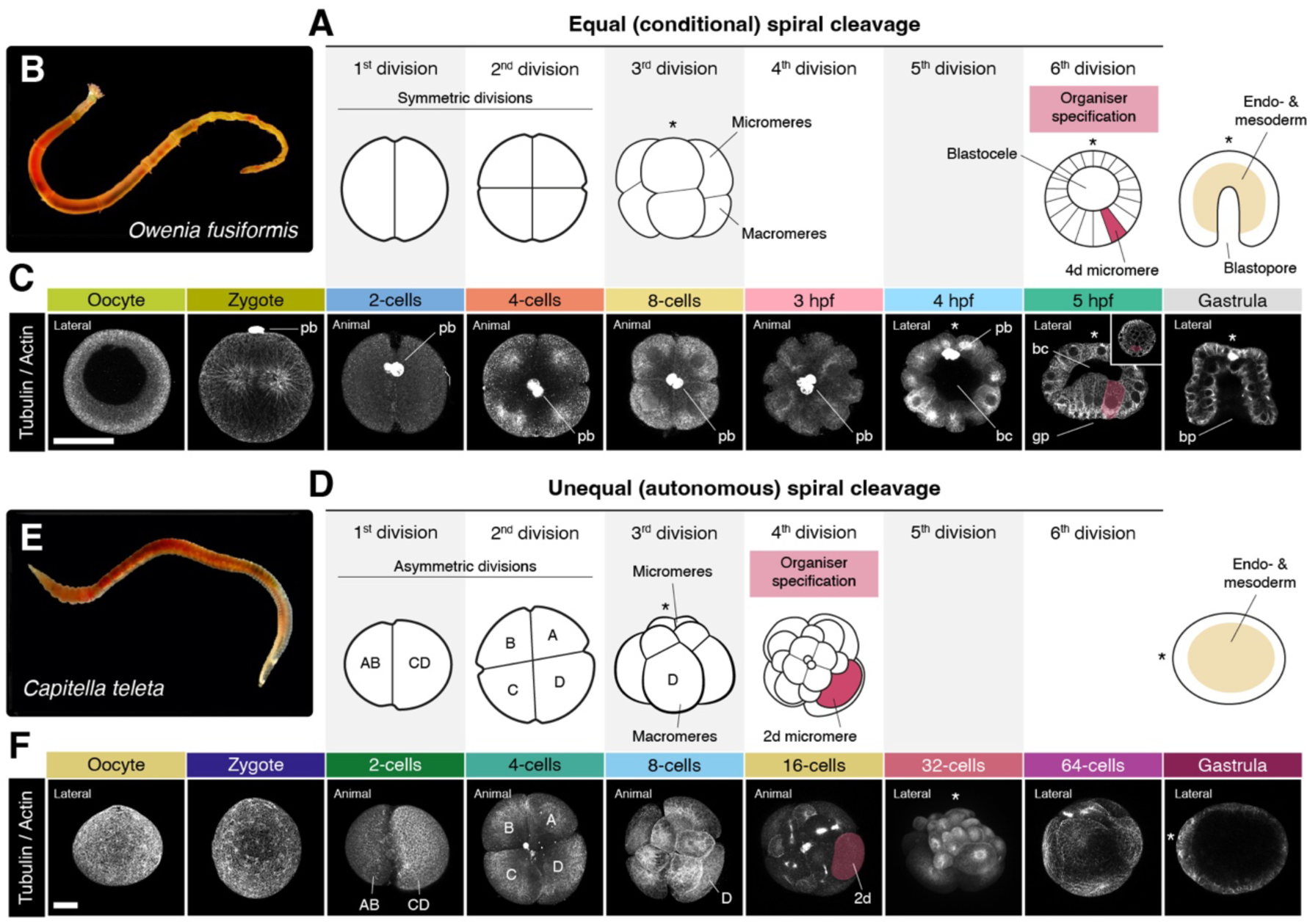
Equal and unequal spiral cleavage in two annelid species. (**A**, **D**) Schematic drawings of equal (or conditional) (**A**) and unequal (or autonomous) (**D**) spiral cleavage as exemplified by the early development of the annelids *O. fusiformis* (**B**) and *C. teleta* (**E**). The drawings highlight the two different types of first zygotic divisions, the idiosyncratic spiral arrangement of the animal and vegetal blastomeres at the 8-cell stage, the different timings of embryonic organiser specification, and distinct gastrula morphologies in embryos that will develop into planktotrophic (*O. fusiformis*) and lecithotrophic (*C. teleta*) larvae. (**C**, **F**) Z-projections of confocal stacks of oocytes and embryos at the exact stages collected for transcriptomic profiling in *O. fusiformis* (**C**) and *C. teleta* (**F**). Samples are stained against tubulin and actin (both in grey) to reveal cell membranes and contours. In *C. teleta* (**F**), the blastomere lineages at the 2-, 4-and 8-cell stages are highlighted. The cell acting as an embryonic organiser (the 4d micromere in *O. fusiformis* and the 2d blastomere in *C. teleta*) is false-coloured in red at the 5 hpf and 16-cell stage, respectively. Drawings are not to scale, and the asterisks indicate the animal/anterior pole. bc, blastocoele; bp, blastopore; hpf, hours post-fertilisation; pb, polar bodies; In (**C**, **F**), scale bars are 50 μm.

Here, we investigate the genome-wide impact of distinct cell fate specification strategies on spiral cleavage. Does the conservation of an ancient programme of cell division patterns and cell lineages constrain the unfolding of developmental programmes during spiral cleavage, or do differences in cell fate specification modes play a more prominent role in defining gene expression dynamics? To answer these, we generated high-resolution transcriptomic profiles at equivalent time points during the early embryogenesis of *Owenia fusiformis* and *Capitella teleta*, two annelid worms with equal/conditional and unequal/autonomous spiral cleavage, respectively (Figure 1) (16, 21, 22). Our findings reveal that these two annelids follow roughly similar transcriptomic transitional phases during spiral cleavage, with maternal genes decaying before the 8-cell stage and the zygotic genome activation likely occurring at the 16-cell stage. Nonetheless, the genes and temporal dynamics defining some of these phases are markedly different and mirror the timing and mechanistic differences in axial patterning and embryonic organiser specification between *O. fusiformis* and *C. teleta* (16, 22). Despite these differences, the embryos of these annelids exhibit a period of maximal similarity –– at the transcriptomic and molecular patterning level –– at the late cleavage and gastrula stage, suggesting that, unlike previous hypotheses (23–26), spiral-cleaving species exhibit a mid-developmental transition, or phylotypic stage, during embryogenesis. Together, our study uncovers an unexpected transcriptomic diversity between species with an otherwise broadly conserved cleavage program, suggesting that distinct cell-fate specification strategies, rather than the conservation of cleavage patterns and overall cell lineages, underpin the evolution of developmental programmes in Spiralia.

## Results

### Global transcriptional dynamics during spiral cleavage

To investigate the global transcriptional dynamics during annelid spiral cleavage, we generated high-resolution time courses of bulk RNA-seq data for the annelids *O. fusiformis* and *C. teleta*, collecting samples in biological duplicates of active or mature oocytes, zygotes, and at each round of cell division until the gastrula stages (Figure 1C, F). In all cases, replicates correlated highly (Supplementary Figure 1A, B; Supplementary Figure 2A, B), with developmental timing accounting for most of the variance (62.4% and 57.6% for *O. fusiformis* and *C. teleta*, respectively) in both species (Figure 2A, B). However, unlike the activated oocyte of *O. fusiformis*, the oocyte stage of *C. teleta* was markedly transcriptionally distinct from the fertilised zygote (Figure 2B; Supplementary Figure 2A), suggesting that either the collected oocytes were not fully mature or that significant transcriptomic changes occur during mating and fertilisation in this species. Since fertilisation is poorly understood in *C. teleta* and cannot be experimentally controlled (27, 28), we could not discern between those two possibilities. Thus, we discarded the oocyte stage for this species in the downstream analyses.

**Figure 2.**
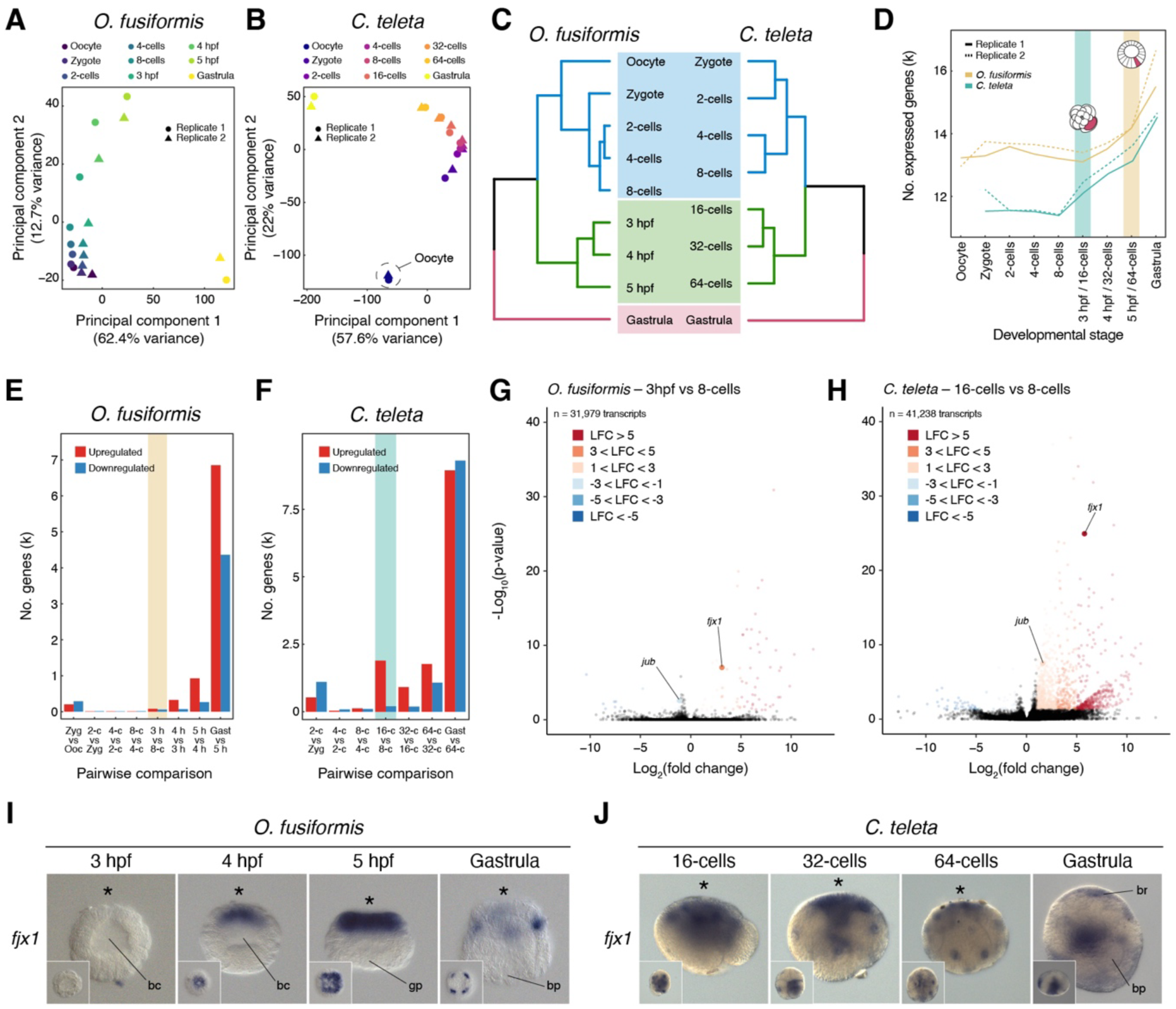
Global transcriptional dynamics during conditional and autonomous spiral cleavage. (**A**, **B**) Principal component analyses support that developmental time accounts for the largest fraction of the variance in the transcriptomic time courses during spiral cleavage in *O. fusiformis* and *C. teleta*. (**C**) Similarity clustering reveals three main sample groups during spiral cleavage in the two species. (**D**) *O. fusiformis* and *C. teleta* show different dynamics in the total number of expressed genes during spiral cleavage, which align with the different timings of organiser specification in these species (highlighted by a yellow and green bar, respectively). (**E**, **F**) Bar plots depicting the number of differentially expressed genes (upregulated in red and downregulated in blue) during spiral cleavage in *O. fusiformis* (**E**) and *C. teleta* (**F**). The proposed timing of zygotic genome activation is highlighted with a yellow and green bar, respectively. (**G**, **H**) Volcano plots indicating up-and downregulated genes in the 8-cell to 16-cell transition in *O. fusiformis* (**G**) and *C. teleta* (**H**). There are only two one-to-one orthologs (*fjx1* and *jub*) that are differentially expressed at this stage in both species. LFC stands for log-fold change. (**I**, **J**) Whole-mount *in situ* hybridisation of *jbx1* from the 16-cell stage (or 3 hpf in *O. fusiformis*) until gastrulation in *O. fusiformis* (**I**) and *C. teleta* (**J**). In *O. fusiformis*, *jbx1* is asymmetrically expressed in the anterior ectoderm. In *C. teleta*, *jbx1* is broadly expressed during cleavage and asymmetrically localised in the anterior neuroectoderm and endoderm in the gastrula.

Similarity clustering revealed three transcriptionally distinct groups during spiral cleavage in the two annelids (Figure 2C). The earliest cluster comprises the oocyte (in *O. fusiformis*) and early cleavage stages up to the 8-cell stage. The late cleavage time points (3 to 5 hours post fertilisation [hpf] in *O. fusiformis* and the 16-cell to 64-cell stages in *C. teleta*) and the gastrula stages make the two other groups. Accordingly, the number of expressed genes (those with transcripts per million [TPM] values above 2; Supplementary Figure 1C; Supplementary Figure 2C) is relatively constant between stages in the first cluster in both species (Figure 2D). However, there is a continuous increase in the number of expressed genes during the late cleavage and, particularly, with gastrulation. This increase in *C. teleta* starts as early as the 16-cell stage and occurs more sharply than in *O. fusiformis*, in which the rise is gradual and not obvious until the 64-cell stage (Figure 2D). Notably, these distinct dynamics mirror the different timings of axial specification in *O. fusiformis* and *C. teleta* (Figure 1A, D; Figure 2C) (16, 22), suggesting that conditional and autonomous specification of the embryonic organiser influence global transcriptional trends during cleavage in these annelids.

Statistically significant changes in gene expression support the distinct transcriptional activation dynamics between *O. fusiformis* and *C. teleta* (Figure 2E, F). While there is a gradual increase in the differentially expressed genes (DEGs) becoming upregulated from 3 hpf onwards in *O. fusiformis*, the number of upregulated DEGs at the equivalent 16-cell stage is more prominent, and a steep change from the levels of DEGs in the previous pairwise comparison in *C. teleta* (78 in *O. fusiformis* vs 1,890 in *C. teleta*) (Supplementary Tables 1– 31). This agrees with the timing of specification of the embryonic organiser in *C. teleta* and supports its role in triggering transcriptional programmes involved in embryonic patterning and cell-fate commitment at that stage in this species (22). In both species, however, gastrulation involves a prominent transcriptional reshaping, with five to seven times more up-and downregulated DEGs at that stage compared with previous cleavage stages (Figure 2E, F). Accordingly, Gene Ontology (GO) terms enriched in the genes upregulated in the gastrula of both species are involved in development and transcriptional regulation (Supplementary Figures 3H, 5G). Interestingly, we identified DEGs between the oocyte and zygote (in *O. fusiformis*) and the 2-cell stage and zygote (in *C. teleta*). In the two annelids, downregulated DEGs are more abundant at these stages (290 in *O. fusiformis* and 1,101 in *C. teleta*), are involved in metabolism and translation (*O. fusiformis*) and protein transport and localisation (*C. teleta*) and might represent maternal genes of quick decay (Figure 2E, F; Supplementary Figures 4A, 6A). Nonetheless, we also detected 203 and 527 upregulated genes in *O. fusiformis* and *C. teleta*, respectively, at those stages (Figure 2E, F). These are enriched for GO terms related to cellular organisation (Supplementary Figures 3A, 5A) and might be differentially polyadenylated transcripts upon fertilisation and the first zygotic division, as observed in other invertebrates (29, 30). Altogether, our high-resolution dataset demonstrates that annelid embryos shared similar transcriptomic phases during spiral cleavage. Yet, the modes and timing of organiser specification influence the span and intensity of these phases during early development.

### Different maternal to zygotic transitions in spiral-cleaving embryos

In *O. fusiformis* and *C. teleta*, the first prominent increase in DEGs occurs at the 16-cell stage, the 4^th^ round of cell division after fertilisation (Figure 2E, F). This suggests that zygotic genome activation, and thus the maternal to zygotic transition (MZT), might occur at that stage, yet with different intensities, in the two annelids. To explore the genes potentially involved in MZT and how this process compares between these species, we used previously calculated functional annotations and one-to-one orthologs in *O. fusiformis* and *C. teleta* (31). Upregulated genes at this stage are enriched for GO categories related to transcription (Supplementary Figure 3E, 5D). In *O. fusiformis*, these involve over ten unclassified TALE homeoboxes, which are expanded in Spiralia and expressed during early cleavage along the animal-vegetal axis (32), and several Fox genes, including many of the expanded *foxQ2* genes that are asymmetrically expressed in the animal pole at 3 hpf (33) and *foxA*, which is a well-known pioneer factor initiating chromatin opening (34, 35). As expected by the higher number, upregulated genes at the 16-cell stage are diverse and involved in many developmental processes in *C. teleta*. These include, among others, genes directly involved in transcriptional regulation (e.g., transcription elongation factors and histone demethylases), cell-fate specification transcription factors (e.g., Six genes, *otx*, *otp*, Fox genes) and signalling pathway components (e.g., Wnt ligands, Frizzled and FGF receptors and TGF-b modulators) (Supplementary Table 24). However, only two genes upregulated at the 16-cell stage in *C. teleta* had a differentially expressed ortholog at 3 hpf in *O. fusiformis* (Figure 2G, H). One –– *fjx1*, the homolog of *four jointed* in *Drosophila melanogaster*, involved in conferring positional information as a potential downstream target of the Notch Delta pathway (36, 37) –– was upregulated in both species, but the other one –– *jub*, a gene encoding for AJUBA, a protein involved in many developmental processes, from transcriptional co-repression to the negative regulation of the Hippo signalling (38) –– was downregulated in *O. fusiformis* (Figure 2G). Therefore, although MZT might occur at the same developmental time, the first signs of zygotic genome activation largely involve different genes in *O. fusiformis* and *C. teleta*.

Given the shared upregulation of *fjx1* at the 16-cell stage, we compared the expression of this gene from this time point until gastrulation in *O. fusiformis* and *C. teleta* (Figure 2I, J). Consistent with its modest upregulation (Figure 2G), we did not detect expression for *fjx1* by whole-mount in situ hybridisation at 3 hpf in *O. fusiformis* (Figure 2I). At 4 and 5 hpf, *fjx1* was, however, strongly expressed in the animal-most ectoderm but not in the top micromeres, becoming restricted to four radially symmetrical cell clusters in the animal hemisphere at the gastrula stage (Figure 2I). This pattern is reminiscent of the expression of several *foxQ2* genes during spiral cleavage in *O. fusiformis*, which are already expressed at the 16-cell stage (33). In *C. teleta*, however, *fjx1* showed a broader expression domain from the 16-to the 64-cell stage, and its expression became restricted to the anterior neuroectoderm and endoderm at the gastrula stage (Figure 2J). Therefore, orthologous genes also follow different spatial expression dynamics upon zygotic activation, which occurs asymmetrically throughout the embryo, at least in *O. fusiformis*.

### Gene-specific transcriptional dynamics differ in spiral cleavage

Although spiral cleavage is a highly stereotypical developmental programme (10–14), our findings at the 16-cell stage hint at gene-specific differences between annelid embryos at similar developmental time points (Figure 2G, H). To attain a comparative, genome-wide view of gene expression dynamics between *O. fusiformis* and *C. teleta*, we first used one-to-one orthologs (n = 7,607) to calculate the Jensen-Shannon index of transcriptomic similarity during early development in the two annelids (Figure 3A). Consistently with our observations at the 16-cell stage, transcriptomic similarity is low during cleavage stages but increases in the late blastula and gastrula stages (Figure 3A). Soft *k*-means clustering of all expressed genes during early embryogenesis (31,323 for *O. fusiformis* and 38,662 for *C. teleta*) revealed five and seven clusters of temporally coexpressed genes in *O. fusiformis* and *C. teleta*, respectively (Figure 3B, C; Supplementary Tables 32, 33). The first two temporally active clusters in both species are maternal genes that are either oocyte-/zygote-specific (cluster 1; 3,593 genes in *O. fusiformis* and 2,934 in *C. teleta*) and have a different codon usage than zygotically-expressed genes (Supplementary Figure 7A, C, D) or that peak shortly after fertilisation (cluster 2; 5,317 genes in *O. fusiformis* and 5,450 in *C. teleta*) (Figure 3B, C). Genes in cluster 1 get quickly removed with fertilisation and the first zygotic division, while genes in cluster 2 decay before the 8-cell stage in both species. Gene Ontology (GO) and KEGG pathway enrichment analyses indicate these clusters are overrepresented with genes involved in metabolism, suggesting an early clearing of transcripts that might be involved in oocyte maturation (Supplementary Figure 8A–F; Supplementary Figure 9A–F). In *O. fusiformis*, 168 of the 290 differentially downregulated genes between the oocyte and the zygote (57.93%; Figure 2D) are oocyte-specific (cluster 1), supporting a rapid elimination of transcripts potentially involved in oocyte maturation upon fertilisation (Supplementary Figure 4A). Likewise, 162 of the 203 upregulated genes with fertilisation (57.93%; Figure 2D) in *O. fusiformis*, and potentially polyadenylated, were not cleared during early cleavage (i.e., were not in clusters 1 and 2), suggesting that, as in other organisms, polyadenylation might be a mechanism to differentially stabilise maternally deposited transcripts (29, 30).

**Figure 3.**
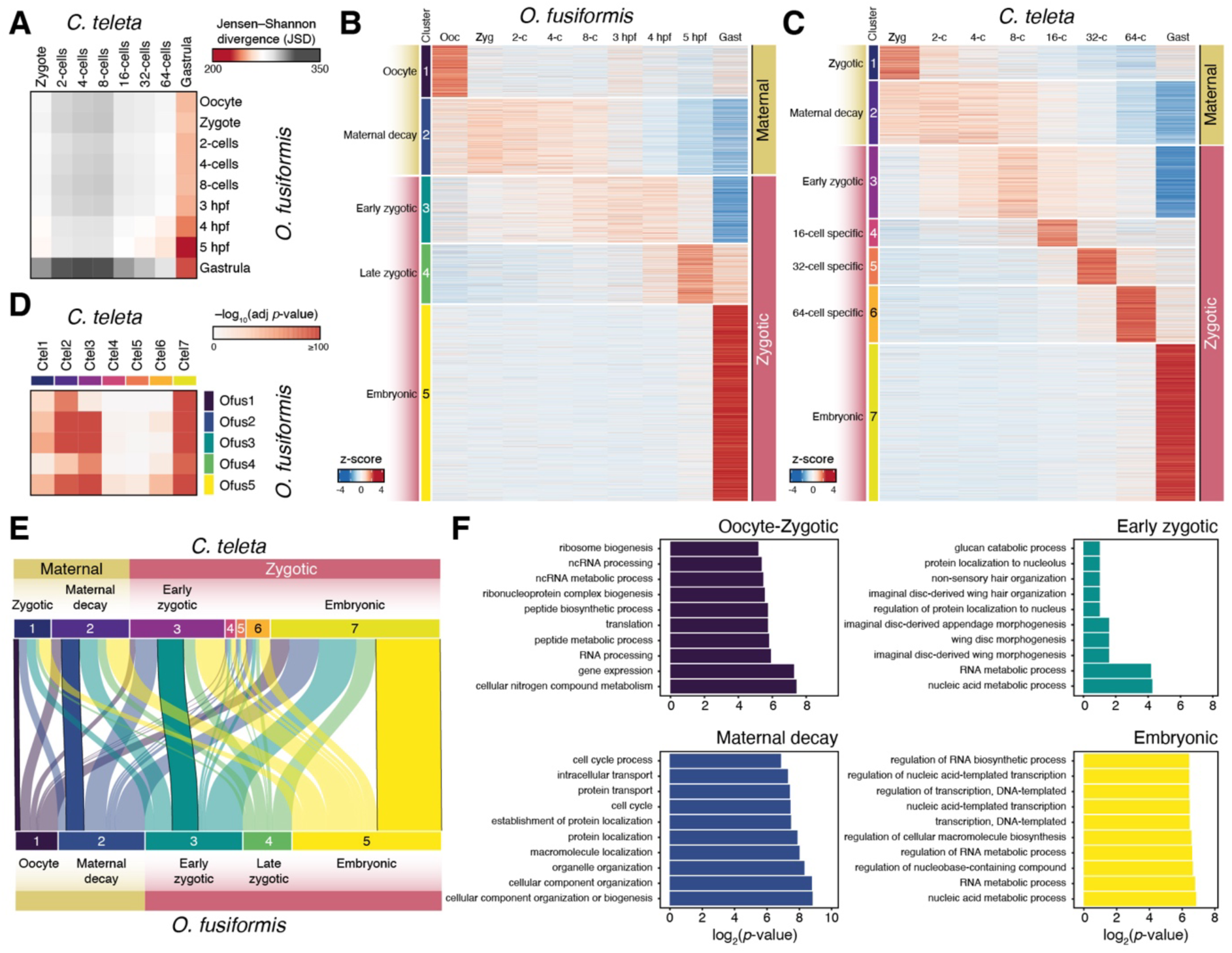
Distinct transcriptomic routes during spiral cleavage in the annelids *O. fusiformis* and *C. teleta*. (**A**) Jensen-Shannon transcriptomic divergence between all possible inter-species pairwise comparisons during the spiral cleavage of *O. fusiformis* and *C. teleta*. The point of maximal transcriptomic similarity between these annelids occurs at the late cleavage and gastrulation. (**B**, **C**) Soft *k*-means clustering of temporally coexpressed genes in *O. fusiformis* (**B**) and *C. teleta* (**C**). In both species, the first two clusters likely represent maternal transcripts of early decay, while the rest largely or entirely comprise zygotically expressed genes. Unlike *O. fusiformis*, *C. teleta* has cleavage-specific clusters at the 16-cell, 32-cell and 64-cell stages. (**D**) Comparison of gene family cluster composition between *O. fusiformis* and *C. teleta*. Although gene-specific transcriptional dynamics are dissimilar, gene family deployment in maternal, early zygotic and embryonic clusters are similar between these annelids. (**E**) Alluvial plot depicting the comparative deployment of one-to-one orthologs (n= 7,607) in clusters of temporally coexpressed genes during spiral cleavage in *O. fusiformis* and *C. teleta*. The embryonic clusters exhibit the most extensive gene composition conservation between species. (**F**) Gene Ontology enrichment of shared orthologs between oocyte/zygote, maternal decay, early zygotic, and embryonic clusters, according to the GO annotation of *O. fusiformis*. While oocyte/zygotic genes are involved in metabolism, shared orthologs of maternal decay are involved in the cell cycle and cellular organisation, and early zygotic and embryonic genes are involved in transcriptional control and development.

In both species, the remaining clusters (3 to 5 in *O. fusiformis* and 3 to 7 in *C. teleta*) comprise genes that increase in expression around the 8-cell stage or later (Figure 3B, C). In *O. fusiformis*, early zygotic genes (cluster 3; 4,610 genes) become more expressed at the 8-cell stage but are primarily restricted to the 3 and 4 hpf stages. They are still involved in housekeeping functions, such as metabolism, cellular transport, and protein/organelle localisation (Figure 3B; Supplementary Figure 8G–I). In contrast, late zygotic genes (cluster 4; 4,109 genes) peak at 5 hpf and are enriched in GO terms involved in MAPK signalling, amongst other signalling pathways (Supplementary Figure 8J–L), which is consistent with the activation of ERK1/2 at this stage to define the embryonic organiser in *O. fusiformis* (16). Lastly, a large (13,695 genes) gene set enriched in GO terms related to animal development (cluster 5) emerges upon gastrulation in *O. fusiformis* (Figure 3B; Supplementary Figure 8M– O). *Capitella teleta* also shows a gene set (cluster 3; 6,161 genes) of early zygotic expression involved in metabolism, cellular transport, and protein localisation that is lowly expressed in the zygote and gradually increases its expression to peak at the 8-cell stage (Figure 3C; Supplementary Figure 9G–I). Likewise, this species also exhibits a large gene cluster restricted to gastrulation (cluster 7; 13,541 genes; Figure 3C). However, unlike *O. fusiformis*, *C. teleta* exhibits stage-specific clusters of temporally coregulated genes for the 16-cell (2,310 genes), 32-cell (3,104 genes), and 64-cell (4,841 genes) stages (Figure 3C). Genes in these clusters involve diverse biological processes, but DNA metabolism and GPCR-mediated neuropeptide signalling are common (Supplementary Figure 9J–R). However, around half of the genes in these clusters do not have a functional annotation (1,040 genes or 45% in cluster 4; 1,565 or 50.42% in cluster 5; and 2,161 genes or 44.64% in cluster 6), indicating that lineage-restricted genes might play a role at these cleavage stages in *C. teleta*. Altogether, our findings support that *O. fusiformis* and *C. teleta* generally follow similar transcriptional trends. However, *C. teleta* shows more complex and dynamic gene coexpression patterns of zygotically-expressed genes, whereas *O. fusiformis* mainly restricts zygotic expression with and after the specification of the embryonic organiser at 5 hpf.

Despite the shared transcriptional trends between *O. fusiformis* and *C. teleta* (Figure 3B, C), the gene-specific transcriptional similarity is low during spiral cleavage (Figure 3A). This might indicate that either cluster composition is dissimilar or similar genes comprise equivalent clusters but expressed at different levels. To test this, we first performed cross-species pairwise comparisons of orthogroup composition between clusters of temporally coexpressed genes (Figure 3D). Clusters comprising genes of maternal decay (clusters 2), early zygotic expression (clusters 3), and embryonic expression (cluster 5 in *O. fusiformis* and cluster 7 in *C. teleta*) showed high inter-species conservation, supporting these include gene families involved in core, conserved embryonic processes, such as early cell cycle progression, and body patterning (Supplementary Figure 8D–I, M–O; Supplementary Figure 9D–I, S–U). However, *C. teleta*’s cleavage-specific clusters showed little correspondence with any cluster in *O. fusiformis* (Figure 3D). Moreover, embryonic clusters in both species, especially in *C. teleta*, showed similarity with earlier gene sets of coexpressed genes (Figure 3D). Next, we compared the cluster allocation of one-to-one orthologs (Figure 3E). Differences in orthogroup composition are general and profound (Figure 3E; Supplementary Figure 10A). For example, 8.18% (622) of embryonic genes in *O. fusiformis* had maternal expression in *C. teleta*; vice versa, 11.29% (859) of embryonic genes in *C. teleta* were maternal in *O. fusiformis*. However, as expected, embryonic (14.92% in *O. fusiformis*), early zygotic (6.02% in *O. fusiformis*), and maternal decay (4.23% in *O. fusiformis*) clusters show the highest (yet still relatively low) proportions of conserved one-to-one orthologs (Supplementary Figure 10A). These shared orthologs are enriched in GO terms for cell-cycle progression and protein/organelle localisation (maternal decay clusters), morphogenesis (early zygotic) and transcription (embryonic) (Figure 3F; Supplementary Figure 10B–F). Although the long evolutionary distance between *O. fusiformis* and *C. teleta* likely influences the extent to which gene-specific transcriptional dynamics differ during cleavage in these species, our findings indicate extensive temporal differences in gene usage between *O. fusiformis* and *C. teleta*. Therefore, only a small proportion of genes might be essential and evolutionarily conserved to maintain the fundamental properties of spiral cleavage and annelid embryogenesis.

### Temporal shifts in the activation of developmental programmes

Because genes conserved between *O. fusiformis* and *C. teleta* are enriched in GO categories associated with morphogenesis and transcription (Figure 3F), we hypothesised that a fraction of those would be transcription factors (TFs) sustaining conserved developmental programmes in early annelid embryogenesis. We thus first characterised the expression dynamics of TFs in these two annelid species. The number of expressed TFs increases as spiral cleavage proceeds in both species (Figure 4A). However, TFs are especially abundant in the embryonic cluster of temporally coregulated genes (Figure 4B), as expected for this gene set to be involved in transcription and activation of body patterning developmental programmes. As per TF class, zinc fingers of class C2H2 dominate the repertoire of expressed TFs (e.g., 454 and 178 at the gastrula stage and embryonic cluster in *O. fusiformis*, respectively; 235 and 187 at the gastrula stage and embryonic cluster in *C. teleta*, respectively) followed by homeodomain-containing and bHLH genes (Supplementary Tables 34–37). To further understand how similar TF expression dynamics are between species, we compared the TF gene family composition of clusters of coexpressed genes (Figure 4C). As with genome-wide approaches, inter-species similarity in TF expression is the highest in embryonic and maternal decay clusters. However, TF gene family composition was also similar between the embryonic cluster in *O. fusiformis* and the zygotic/maternal decay cluster in *C. teleta* and between the embryonic cluster in *C. teleta* and the maternal decay/early zygotic cluster in *O. fusiformis* (Figure 4C). Therefore, the similarity in gene family usage at equivalent timings of development between these annelids is not only driven by the shared use of structural genes that are essential for animal cleavage (e.g., cell cycle and cytoskeleton-related genes) but also similar transcription factors.

**Figure 4.**
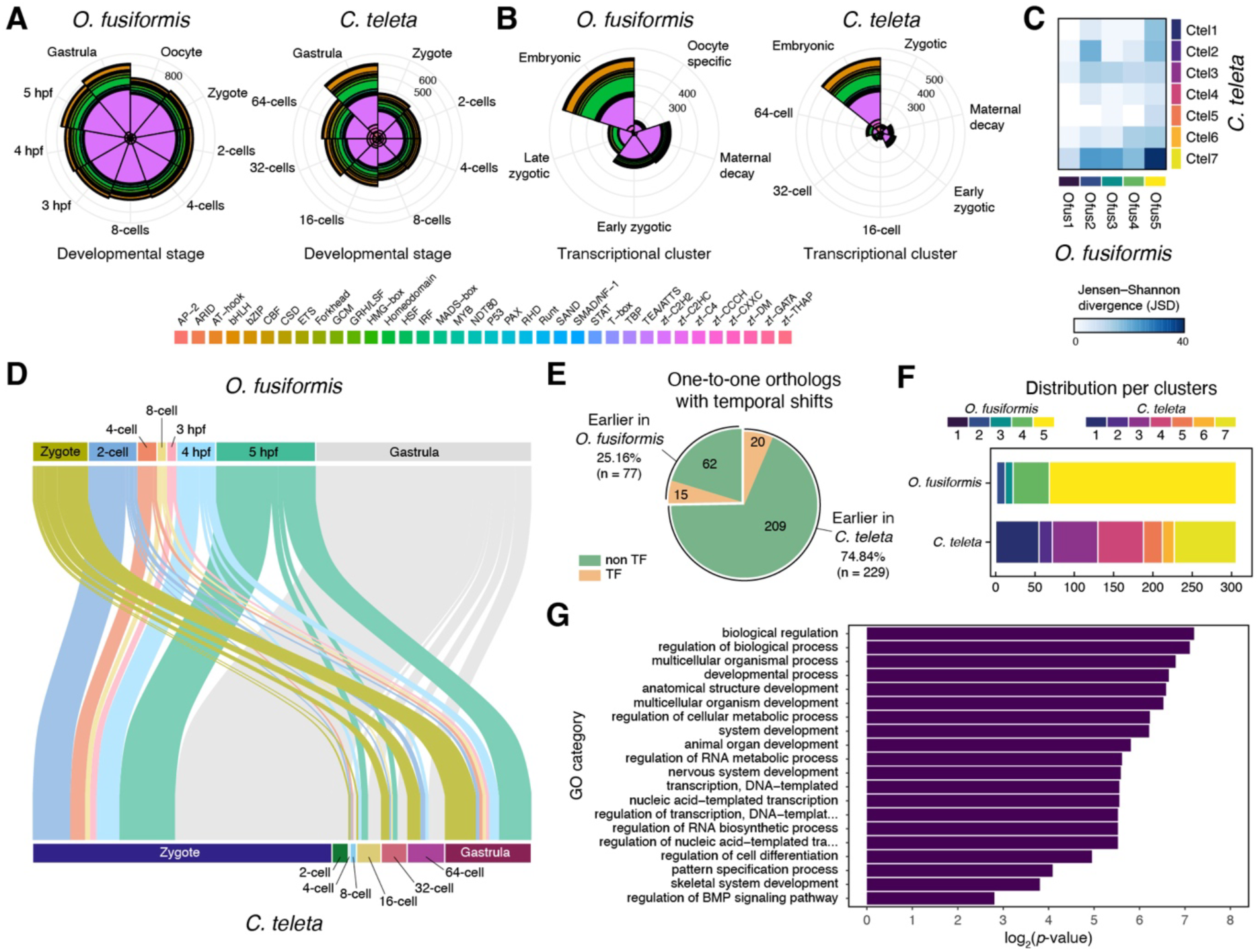
The comparative dynamics of transcription factor activation in conditional and autonomous annelids. (**A**, **B**) Nightingale Rose Charts of transcription factor (TF) distribution according to developmental time points (**A**) and clusters of coexpressed genes (**B**). (**C**) Comparison of gene family cluster composition between *O. fusiformis* and *C. teleta*. TF similarity in clusters of genes showing maternal decay and embryonic expression is high. However, there are prominent shifts in TF gene families allocated to the embryonic cluster in one species to earlier clusters in the other. (**D**) Alluvial plot depicting the comparative deployment of one-to-one orthologs (n= 306) exhibiting shifts in temporal activation (TPM > 2) during spiral cleavage in *O. fusiformis* and *C. teleta*. (**E**) Pie chart showing that most genes with a temporal shift between *O. fusiformis* and *C. teleta* are expressed earlier in the latter. (**F**) The bar plots indicate the allocation of genes with a temporal shift to the clusters of coregulated genes in *O. fusiformis* and *C. teleta*. Most genes exhibiting temporal shifts between these species belong to late clusters in *O. fusiformis* and shift towards earlier clusters in *C. teleta*. (**G**) Gene Ontology enrichment of one-to-one orthologs (n=306) exhibiting temporal shifts in transcriptional activation between *O. fusiformis* and *C. teleta*, according to the GO annotation of *C. teleta*.

Despite the overall similarities in TF usage in early and late cleavage, genome-wide and TF-only inter-species cluster composition comparisons also support differential temporal activation of orthologous gene families during spiral cleavage in *O. fusiformis* and *C. teleta* (Figure 3D; Figure 4C). To identify the exact genes exhibiting these temporal shifts in expression, we compared the activation timing (TPM > 2) of one-to-one orthologs during spiral cleavage in the two annelids (Figure 4D). Consistent with inter-cluster comparisons, the majority of orthologs exhibit a shift from gastrula and late spiral cleavage (5 hpf) in *O. fusiformis* to early cleavage stages, including the zygote, in *C. teleta* (Figure 4D). Indeed, temporal shifts in gene expression from late stages in *O. fusiformis* to earlier stages in *C. teleta* are more than twice as common than in the other direction (Figure 4F). Most (75.98%) of the genes –– including 17 out of the 20 TFs (85%) –– exhibiting temporal shifts in *O. fusiformis* belong to clusters 4 (late zygotic, with a peak of expression at 5 hpf) and 5 (embryonic, peaking at gastrulation) (Figure 4E, F; Supplementary Tables 38, 39). However, in *C. teleta*, the genes with heterochronic shifts exhibit more diverse temporal dynamics, with more than half belonging to the early clusters 1 to 4 (zygote-specific, maternal decay, early zygotic and 16-cell specific) (Figure 4F). Generally, the genes exhibiting temporal shifts in transcriptional activation between these species are enriched in GO terms related to development, cell differentiation, transcriptional regulation, and intercellular signalling (e.g., BMP) (Figure 4G). Maternal genes in *O. fusiformis* that are expressed later during cleavage in *C. teleta* are enriched in GO terms related to ERK1/2 regulation, neuronal development, and DNA methylation, among others (Supplementary Figure 12). Inversely, maternal genes in *C. teleta* expressed later in *O. fusiformis* are enriched in GO categories associated with diverse processes, including eye development, the Notch, JNK and MAPK pathways, and cell adhesion and migration (Supplementary Figure 13). Our comparative analyses have thus identified a relatively reduced gene set (306 orthologs, including 35 DNA binding genes) for future functional investigations, as they might be involved in the early developmental differences in organising and cell fate specification between these two annelids.

### Shared expression domains of orthologous TFs in annelid gastrulae

To start to investigate the implications of temporal shifts in TF activation during spiral cleavage in *O. fusiformis* and *C. teleta*, we focused on seven TFs with well-known roles during animal development (*pax2/5/8*, *tbx2/3*, *vsx2*, *AP2*, *uncx*, *HNF4* and *prop1*) (16, 39–55) that follow the most common temporal change between these two annelids (Figure 4D, F), namely expressed at or after organiser specification in *O. fusiformis* (5 hpf and gastrula) but at earlier time points in *C. teleta*. In agreement with the detected expression in the RNA-seq time course (Supplementary Table 40), none of these genes show expression during cleavage until 5 hpf in *O. fusiformis* (Figure 5A–G). The only exception is *pax2/5/8*, which, as expected, exhibits broad maternal expression throughout the embryo (Figure 5A). At 5 hpf, however, all TFs show restricted expression domains (Figure 5A–G). *pax2/5/8* is expressed in a few apical ectodermal cells that extend to encircle the entire anterior apical ectoderm in the gastrula (Figure 5A). Additionally, *pax2/5/8* is detected on one side, potentially the posterior, of the blastopore in *O. fusiformis* (Figure 5A). As with *pax2/5/8*, *tbx2/3*, *vsx2*, and *uncx* are also detected in the apical ectoderm at 5 hpf and gastrula stages in *O. fusiformis*, but also in the gastral plate, endoderm, and the posterior blastopore rim for *uncx* (Figure 5B–D). The TF *AP2* show expression in discrete ectodermal cells in the future dorsoposterior side of the embryo (Figure 5E). Finally, *HNF4* and *prop1* are expressed in the gastral plate at 5 hpf (Figure 5F, G). While we did not detect expression of *HNF4* at the gastrula stage for *O. fusiformis*, *prop1* is expressed in the internalised cells and seven further bilaterally symmetrical clusters of two cells around the equatorial ectoderm of the gastrula (Figure 5F, G).

**Figure 5.**
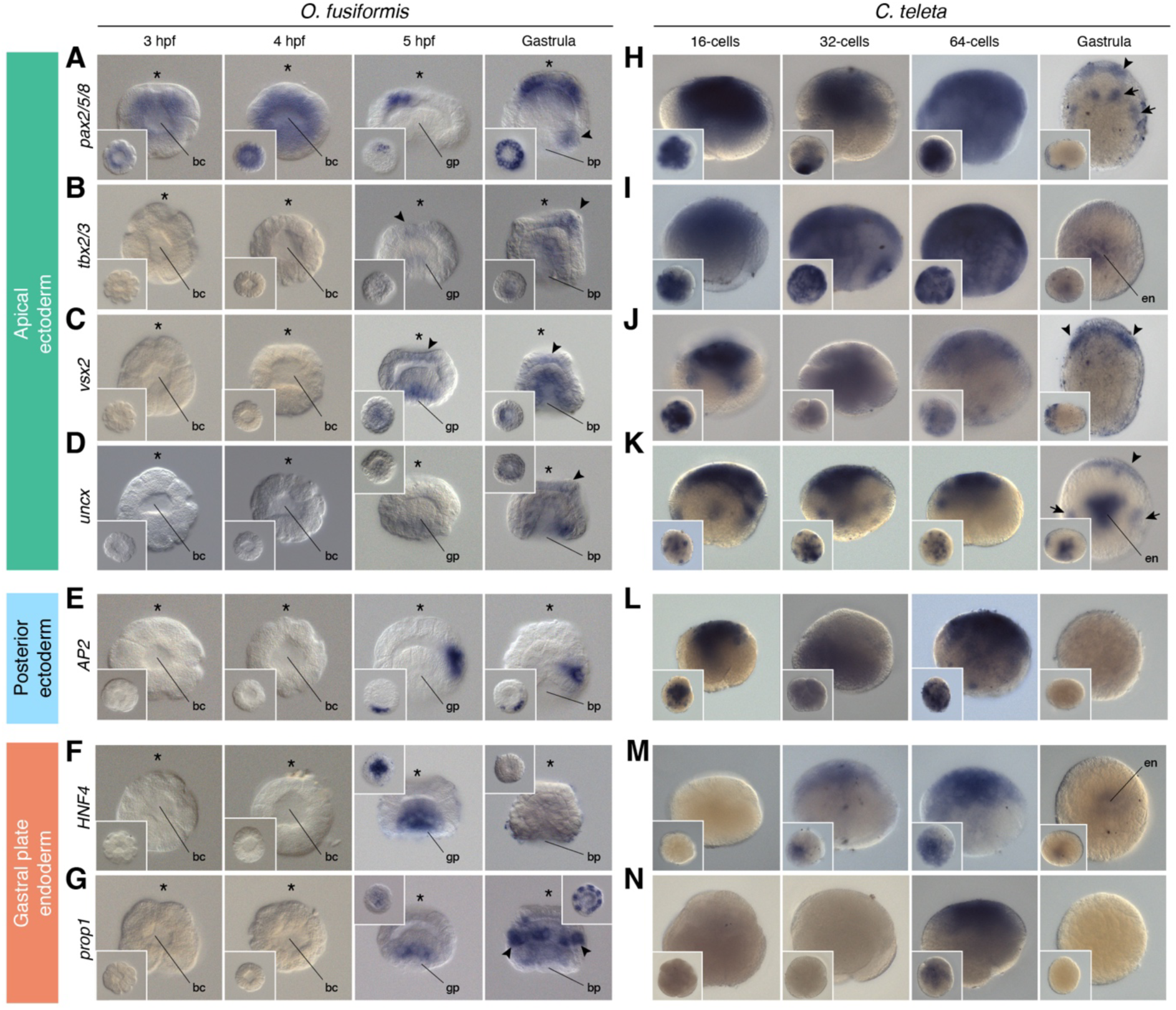
Spatial expression dynamics of orthologous transcription factors in *O. fusiformis* and *C. teleta*. (**A**–**N**) Whole-mount *in situ* hybridisation of seven orthologous transcription factors (*pax2/5/8*, *tbx2/3*, *vsx2*, *uncx*, *AP2*, *HNF4*, and *prop1*) during mid and late cleavage and at the gastrula stage in *O. fusiformis* (**A**–**G**) and *C. teleta* (**H**–**N**). In *O. fusiformis*, *pax2/5/8* (**A**), *tbx2/3* (**B**), *vsx2* (**C**) and *uncx* (**D**) are expressed in the anterior neuroectoderm (arrowheads), as well as in the posterior blastoporal rim (*pax2/5/8* and *uncx*, arrows) and endoderm (*tbx2/3* and *vsx2*). *AP2* (**E**) is expressed in the posterior ectoderm (arrowhead), and *HNF4* (**F**) and *prop1* (**G**) are in the gastral plate before gastrulation and in seven equatorial clusters of two ectodermal cells in the gastrula (*prop1*, arrowheads). No expression is detected before 5 hpf for any of these genes. In *C. teleta*, *pax2/5/8* (**H**), *tbx2/3* (**I**), *vsx2* (**J**), *uncx* (**K**) and *AP2* (**L**) are broadly expressed during mid and late cleavage. In the gastrula, *pax2/5/8*, *tbx2/3*, *vsx2* and *uncx* are detected in the anterior neuroectoderm (arrowheads), four cell clusters posterior to the foregut (*pax2/5/8*, arrows), the endoderm (*tbx2/3* and *uncx*) and two lateral mesodermal clusters (*uncx*, arrows). *HNF4* (**M**) is broadly detected at 32-and 64-cells and in the endoderm of the gastrula, and *prop1* (**N**) expression is only apparent at the 64-cell stage. Insets are animal/vegetal views, except for the gastrula stage in **H**–**N**, which are lateral views.

Consistent with in silico datasets (Supplementary Table 40), these seven TFs were expressed during mid and late cleavage in *C. teleta* (Figure 5H–N). However, unlike in *O. fusiformis*, the early expression of these TFs in *C. teleta* is broad and generally distributed throughout all embryonic blastomeres for most of the genes, and more restricted expression domains only appeared at the gastrula stage (Figure 5H–N). At that stage, *pax2/5/8* is detected in the anterior ectoderm (i.e., the developing brain) and four ventral clusters posterior to the foregut (Figure 5H). Similarly, *tbx2/3*, *vsx2* and *uncx* are also expressed in the anterior neuroectoderm of the gastrula. Additionally, *tbx2/3* and *uncx* show expression in the internalised endoderm and *uncx* is also detected in two lateral domains, reminiscent of the most lateral clusters of *pax2/5/8* that might correspond to the developing mesoderm (Figure 5I, J). Finally, we did not observe expression for *AP2* and *prop1* in the gastrula, but *HNF4* is weakly expressed in the endoderm of *C. teleta* at that stage (Figure 5L–N). Altogether, the comparison of the expression domains of these seven TFs is consistent with the higher transcriptional similarity at the gastrula stage between *O. fusiformis* and *C. teleta* (Figure 3A), as most of these genes (*pax2/5/8*, *tbx2/3*, *vsx2*, *uncx*, and *HNF4*) have shared expression patterns in the late blastula and gastrula despite their different transcriptional dynamics during early cleavage.

### An hourglass-like dynamic of gene expression in spiral cleavage

Our findings support the gastrula as a generally conserved developmental stage –– in transcriptional levels, orthogroup and gene deployment, and gene expression patterns –– during early annelid embryogenesis despite the diverse morphologies at this embryonic period. Indeed, the comparative analysis of transcriptomic similarity throughout the entire development of *O. fusiformis* and *C. teleta* (from oocyte to competent larvae and juvenile stages) strengthens that the late cleavage and gastrulation stages are the most transcriptionally similar stages in the life cycle of these species, followed by the juvenile stages (Figure 6A). The late cleavage and gastrulation thus act as a mid-developmental transition between two phases of higher transcriptomic dissimilarity, namely an earlier period during spiral cleavage and a later phase during larval development. Notably, when comparing annelids with *Crassostrea gigas*, a bivalve mollusc with autonomous spiral cleavage, the late spiral cleavage and gastrulation are not periods of maximal transcriptomic similarity, but spiral cleavage is still a phase of high transcriptional dissimilarity (Figure 6B; Supplementary Figure 14). Instead, the larval stages act as stages of maximal transcriptomic similarity, consistent with previous comparisons amongst metazoan larvae (31).

**Figure 6.**
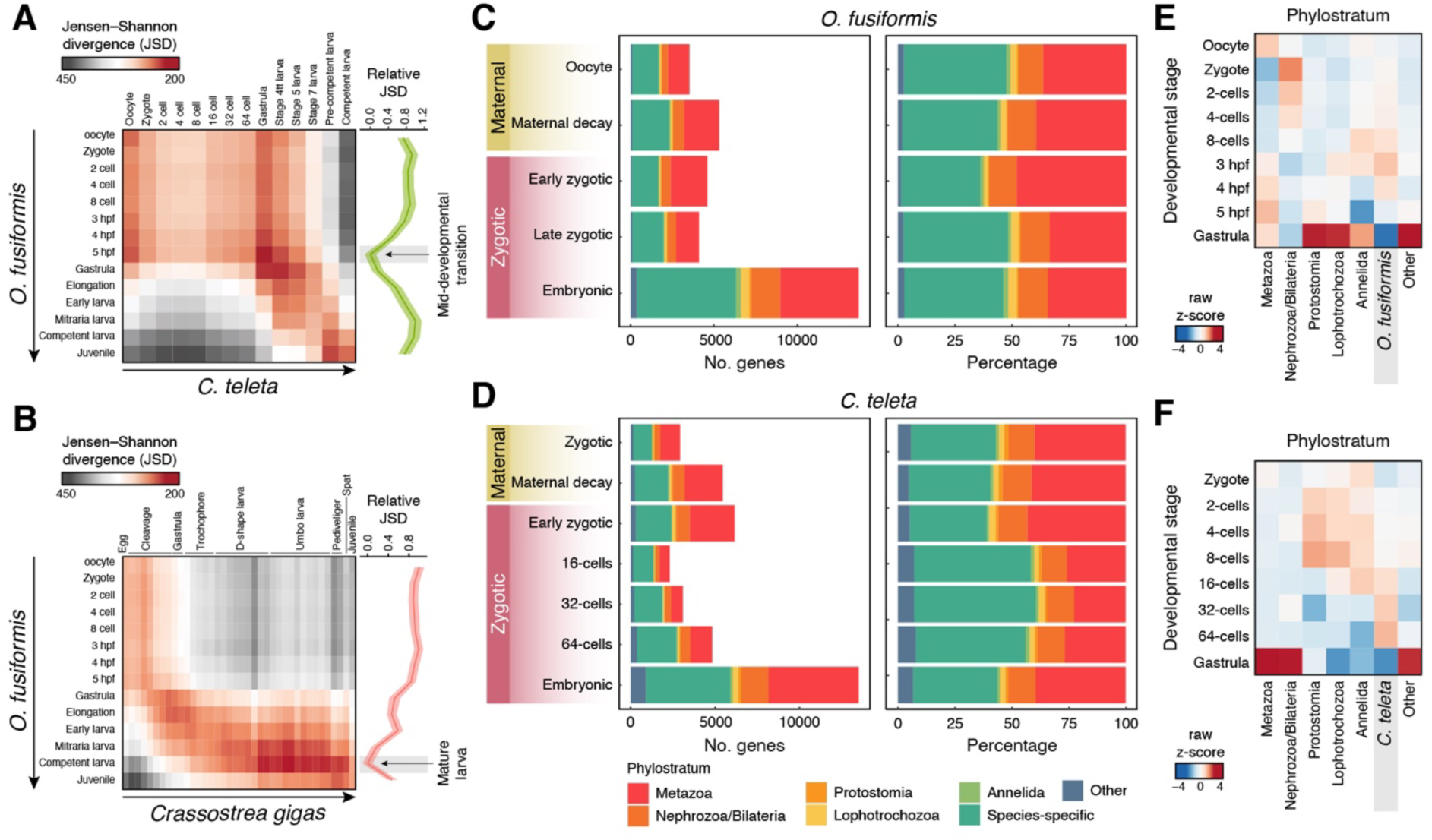
A mid-developmental transition in spiral cleavage. (**A**, **B**) Jensen-Shannon transcriptomic divergence between all possible inter-species pairwise comparisons during the entire life cycle, from oocyte to juvenile or competent larva of *O. fusiformis* and *C. teleta* (**A**) and *O. fusiformis* and *Crassostrea gigas* (**B**). The point of maximal transcriptomic similarity between these annelids occurs at the late cleavage and gastrulation, while it happens at the ciliated larval stage between *O. fusiformis* and *C. gigas*. (**C**, **D**) Bar plots indicate the distribution of expressed genes according to their age or phylostratum at each cluster of temporally coregulated genes in *O. fusiformis* (**C**) and *C. teleta* (**D**). More ancestral genes represent around half of the transcriptome during spiral cleavage in both species, except in the 16-cell, 32-cell and 64-cell stage-specific clusters of *C. teleta*. (**E**, **F**) The heatmaps depict the stages with the highest expression of genes according to age. In *O. fusiformis* (**E**), metazoan, protostomian and lophotrochozoan genes are more expressed at the 5 hpf and gastrula, while in *C. teleta* (**F**), metazoan and bilaterian are highly expressed at the gastrula stage.

Deployment and enrichment in genes of ancestral origin have been associated with developmental stages of high inter-species conservation (26, 56). To assess this in annelids, we assigned genes based on their phylogenetic origin and evaluated their contribution to the clusters of coregulated genes. In both species, old genes (i.e., ancestral to Metazoa and originated in Bilateria, Protostomia and Spiralia) account for more than 50% of the genes in all but the oocyte-specific clusters in *O. fusiformis* and all but the cleavage-specific clusters (16-cell, 32-cell, and 64-cell) in *C. teleta* (Figure 6C, D). These cleavage-specific clusters exhibit more than 50% of *C. teleta*-specific genes (Figure 6D), supporting the lack of functional annotation for many of the genes expressed at those time points in this annelid. Consistent with the gene distribution by phylostratum, the expression of genes of Metazoa, Protostomia, and Spiralia origin is higher at the gastrula stage in *O. fusiformis* (Figure 6E), and Metazoa/Bilateria genes are highly expressed in the gastrula of *C. teleta* (Figure 6F). However, the expression of species-specific genes is higher during mid-cleavage stages in both annelids, from the 8-cell stage to 4 hpf in *O. fusiformis* and from the 16-cell to the 64-cell stages in *C. teleta* (Figure 6E, F). Therefore, our findings indicate that annelids undergo a mid-developmental transition in their life cycle around gastrulation when embryos deploy more ancestral genes and exhibit a high degree of transcriptomic and gene expression pattern similarity.

## Discussion

Our work profiles, with an unprecedented temporal resolution for spiral cleavage (57–60), the transcriptomic dynamics during early embryogenesis in *O. fusiformis* and *C. teleta*, two annelids with equal/conditional and unequal/autonomous spiral cleavage, respectively. Although the evolutionary distance between *O. fusiformis* and *C. teleta* likely influences some of our observations, our extensive dataset defines a novel comparative framework to investigate the genetics and mechanisms controlling and diversifying spiral cleavage, one of the most ancient and broadly conserved cleavage programmes (10–12).

### The early transcriptomic dynamics in spiral cleavage

The transition from a maternally regulated development to one controlled by the zygotic genome –– the so-called maternal to zygotic transition –– is a fundamental event in early animal embryogenesis (61–64). This involves the degradation of maternal transcripts and proteins and the epigenetic remodelling of the zygotic genome to enable transcription (i.e., the zygotic genome activation). These often, but not always, coincide with the desynchronisation of cell cycles between different embryonic regions (the “mid-blastula transition”) (65–67). Notably, the timing of these events varies dramatically, even between phylogenetically closely related lineages (63, 67). Yet, how and when these critical steps occur during embryogenesis remains unknown in most animals, especially those exhibiting spiral cleavage. In the annelid leech *Helobdella triserialis*, which exhibits a modified spiral cleavage (68), embryonic transcription begins early, when the embryo has ∼20 cells, only in a subset of blastomeres, and is essential to control spindle orientation (69). Based on transcriptomic profiling, zygotic genome activation might occur around the 64-cell stage in the annelid *Platynereis dumerilii* (59). However, only two earlier time points (the 8-and 30-cell stages) were sampled in that study. In *O. fusiformis* and *C. teleta*, maternal transcripts are largely cleared in two waves by the 8-cell stage, and the first signs of gene upregulation occur immediately after, at the 16-cell stage (the 4^th^ cell cycle) in both species (Figure 2D–F; Figure 7A). Notably, the first cleavage asymmetry in *O. fusiformis* occurs two cell cycles later, with the formation of the 4q micromeres, one of which will become the embryonic organiser (16). Therefore, the maternal to zygotic transition might occur around the 16-cell stage and not coincide with the mid-blastula transition in annelids, yet inter-species variation is likely. A more extensive taxon sampling that includes other groups with spiral cleavage and more precise approaches to measuring zygotic transcription than poly(A) RNA-seq (61, 63) will provide a more comprehensive view of this essential developmental event in Spiralia.

**Figure 7.**
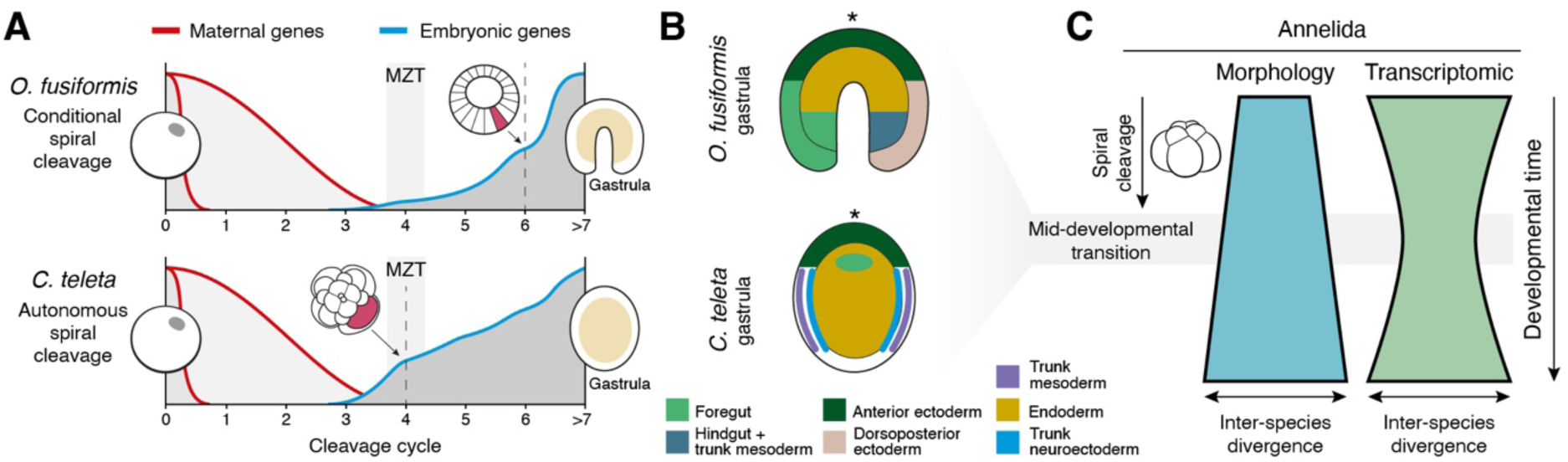
Transcriptional and morphological dynamics are decoupled in spiral cleavage. (**A**) Schematic summary of the transcriptomic dynamics during spiral cleavage in *O. fusiformis* and *C. teleta*. Two waves of mRNA decay (one that is oocyte/zygote-specific and another that extends to about the 8-cell stage) occur during early cleavage. Zygotic genome activation and the maternal to zygotic transition (MZT) likely occurs at the 4^th^ cell division (the 16-cell stage) and more intensely in *C. teleta*, as it coincides with the specification of the embryonic organiser (schematic drawing and dotted vertical line). The large increase in zygotic transcriptional activity occurs at the 6^th^ cell division (64-cell stage) in *O. fusiformis*, when the embryonic organiser is established in this species (dotted line and schematic drawing). (**B**) Schematic comparison of broad areas of developmental competence at the gastrula stage between *O. fusiformis* and *C. teleta* (see main text for details). (**C**) Differently from other animal phyla, morphological and transcriptomic similarities are uncoupled during development in Annelida. The early embryonic phase of spiral cleavage is morphologically stereotypical but transcriptionally divergent. However, annelid embryos converge into a broad transcriptomic and molecular patterning similarity phase by the gastrula stage, which acts as a mid-developmental transitional period. Annelid embryos diverge (morphologically and transcriptionally) upon gastrulation as they develop into lineage-specific larval forms. Drawings are not to scale.

Although it happens at similar developmental time points, the zygotic genome activation is more pronounced in *C. teleta* than in *O. fusiformis* at the 16-cell stage (Figure 2E, F; Figure 7A). This coincides and is consistent with the specification of the embryonic organiser and the activation of head patterning programmes at this stage in *C. teleta* (Figure 1D) (22, 70). Similarly, the specification of the embryonic organiser by the 64-cell stage is concomitant with an increase in gene upregulation in *O. fusiformis* (Figure 2E) (16). As observed in the leech *H. triseralis* (69), our data also support that zygotic genome activation is not uniform throughout the embryo, at least in *O. fusiformis*, as some of the upregulated genes, such as the expanded *foxQ2* genes (33) and *fjx1*, show asymmetric expression along the animal-vegetal axis at these early stages of development. These markers concentrate in the animal ectodermal pole (33) (Figure 2I), which generates anterior neuroectodermal derivatives in most spiralians and bilaterians (13, 14, 71). This reinforces the hypothesis that anterior neural structures develop early and more autonomously in embryos with both autonomous and conditional modes of spiral cleavage (70). In equal/conditional spiral cleavage, the posterior/vegetal pole likely remains uncommitted as new tiers of micromeres are produced (72, 73) and until the inductive signalling event mediated by the FGF-ERK1/2 pathway primes one of the blastomeres to become the embryonic organiser (16–18), restricting anterior fates and activating posterodorsal developmental on one side of the embryo (16). Therefore, although embryos with equal/conditional and unequal/autonomous spiral cleavage share general transcriptional dynamics during early embryogenesis (Figure 7A), the different modes of cell-fate specification, which determine the timing of axial patterning and the onset of body regionalisation, outweigh the conservation of the cleavage program and cell lineages in shaping specific transcriptional trends during early embryogenesis.

### The transcriptomic convergence during axial patterning in annelid embryos

Despite their different early transcriptional dynamics and modes of cell fate specification, late cleavage and gastrula stages are the most similar between *O. fusiformis* and *C. teleta*. The analysis of seven transcription factors activated late (5 hpf and gastrula) in *O. fusiformis* but much earlier in *C. teleta* demonstrates that similarity in transcription levels at the gastrula stage is often accompanied by similarity in expression domains (Figure 5). This is consistent with the study in these annelids of candidate transcription factors with evolutionarily conserved expression domains across distantly related animals that act as ‘marker’ genes of the development of specific structures. Foregut genes, such as *nkx2.1*, *foxA* and *gsc*, are expressed in the prospective oral ectoderm of the gastrula, while endodermal markers, such as *GATA4/5/6* paralogs, localise to the internalised cells (Figure 7B) (74–76). Likewise, anterior neuroectodermal genes (e.g., *sox* genes and those in this study) are expressed in the animal/anterior pole in the gastrulae of *O. fusiformis* and *C. teleta* (Figure 7B) (77–79). Notably, however, the two species differ in the extent and levels of trunk mesodermal and neuroectodermal gene expression (e.g., *pax3/7*, *evx*, *cdx*, *twist*). While the precursors of the ecto-and mesodermal trunk occupy a large lateral domain in the gastrula of *C. teleta* (80–82), these are restricted to one side of the blastoporal rim and the few possible descendants of the embryonic organiser in *O. fusiformis* (16, 74, 83) (Figure 7B). These differences reflect the temporal and developmental differences in the trunk formation in these two species, ultimately linked to their distinct larval types and life cycle strategies (31). Equally, in annelids with heteromorphic larval types (i.e., planktotrophic and lecithotrophic), such as *Streblospio benedicti*, genes exhibiting temporal shifts are connected to morph-specific developmental and physiological traits, like the formation of a functional gut in the planktotrophic type and the metabolism of the maternal nourishment in the lecithotrophic (58). Therefore, despite the conservation of the cleavage programme and the early transcriptional differences, a shared body and molecular patterning that reflects subsequent developmental species-specific traits only emerge during gastrulation in annelid embryogenesis.

### A mid-developmental transition in annelid development

In some animal clades, there is a period in mid-development when phylogenetically related taxa exhibit a significant degree of morphological and transcriptomic similarity (84–86). This is thought to be the stage at which the body plan’s essential traits of the clade are established, and thus, this phase is also termed the phylotypic stage (87, 88). Although the phylotypic stage concept is contested (89–92), an equivalent phenomenon has been described during plant, fungal, and brown algal development (93–95). Therefore, the phylotypic stage might reflect an emergent property of multicellular systems that occurs when genetic programs become active and interact to define the different body regions, creating a stage more sensitive to evolutionary change (96–103). From a morphological standpoint, however, the highest degree of embryonic similarity amongst the disparate and distantly related animal groups with spiral cleavage occurs during the earliest zygotic divisions (Figure 7C). This has led some authors to claim there is no phylotypic stage in Spiralia (23), while others have argued the larva is the phylotypic stage in annelids and molluscs (24–26, 104). Yet, life cycles are vastly diverse in Spiralia, and larvae differ dramatically in their morphology and ecology, even within a phylum (105, 106). Thus, a mid-developmental phylotypic stage in Spiralia has remained controversial.

Our data highlights the gastrula as a critical developmental time point in spiral cleavage (Figure 7B, C). Although the amount of maternally provided nourishment heavily influences embryonic morphologies and morphogenetic processes at this stage (e.g., gastrulation by invagination or epiboly), the gastrulae of *O. fusiformis* and *C. teleta* exhibit high overall transcriptomic similarity, shared molecular expression patterns, and areas of developmental competence (Figure 7B; see above). Morphological and transcriptomic similarities are thus uncoupled during spiral cleavage, indicating that the extreme conservation of this early embryogenesis relies on a limited gene set and/or maternal proteins contributing to conserved cellular physical mechanics (107). Moreover, developmental system drift in the specification of homologous cell types and body regions might be widespread in these lineages, even between relatively closely related species (108). Therefore, unlike in other animal groups like vertebrates and insects, a mid-developmental phylotypic stage around gastrulation only occurs at the transcriptomic level in Spiralia. This stage sits at the transition from an early morphologically similar but transcriptionally divergent phase of cleavage to a late morphologically and transcriptionally divergent period during organogenesis that results in the formation of a larva or juvenile (Figure 7C). This scenario challenges previous interpretations of spiralian development (23–26, 104), proposing a stage in embryogenesis where the astonishing body plan diversity of the animal lineages with spiral cleavage might emerge.

## Conclusions

Our work provides an unparalleled characterisation of the transcriptional dynamics during early embryogenesis in *O. fusiformis* and *C. teleta*. These two annelids exhibit spiral cleavage but autonomous and conditional specification of their body plans, respectively. Unexpectedly, the transcriptomic profiles during their early development are highly divergent, influenced by their strategies and different timings to define the embryonic organiser and bilateral symmetry. This indicates the high conservation of the spiralian cleavage programme does not constrain the deployment of transcriptional programmes during the development of these annelids. Thus, maternal proteomes and/or biophysical properties might play a more significant role in maintaining these species’ shared spiral cleavage pattern of cell divisions. Yet, the embryos of these two annelids converge in their transcriptomic and body patterning profiles at gastrulation, which appears as a mid-developmental transitional phase in annelid embryogenesis. Altogether, our study provides a unique, high-resolution resource to investigate the early stages of animal embryogenesis in two emerging spiralian models, proposing new conceptual scenarios to explore the developmental mechanisms underpinning the conservation and diversification of spiral cleavage, an iconic and ancient mode of animal development.

## Methods

### Animal culture and sample collection

Sexually mature *Owenia fusiformis* Delle Chiaje, 1844 adults were collected from subtidal waters near the Station Biologique de Roscoff and cultured in the laboratory as previously described (21). In vitro fertilisation and collection of embryonic stages were performed as outlined (21). *Capitella teleta* (Blake, Grassle & Eckelbarger, 2009) was cultured, and embryos were collected following established protocols (109). For oocyte collection, females whose offspring were used to collect cleavage stages were isolated and dissected after one week with abundant mud to obtain their oocytes.

### Library preparation and sequencing

*O. fusiformis* developmental samples encompassing active oocyte, zygote, 2-cell, 4-cell and 8-cell stages, 3 hours post-fertilization (hpf), 4 and 5 hpf (coeloblastula), and gastrula (9 hpf) were collected in duplicates, flash frozen in liquid nitrogen, and stored at –80 °C for total RNA extraction. Samples within replicates were paired, with each one containing 500 embryos coming from the same in vitro fertilisation. Reciprocal developmental stages for *C. teleta*, namely the oocyte, zygote, 2-, 4-, 8-, 16-, 32-, and 64-cell stages, and gastrulae were collected in duplicates using similar genetic pools of different brood tubes. Total RNA was isolated using a Monarch Total RNA Miniprep kit (New England Biolabs) following the supplier’s recommendations and used to prepare strand-specific mRNA Illumina libraries that were sequenced at the Oxford Genomics Centre (University of Oxford, UK) over three lanes of an Illumina NovaSeq6000 system in 2 × 150 bp mode to a depth of around 50 million reads per sample (Supplementary Tables 41, 42).

### RNA-seq analyses

To profile gene expression dynamics during development, we first utilised Kallisto (0.46.2) (110) to map RNA-seq reads to the reference gene models of *O. fusiformis* and *C. teleta*. Mapping statistics are provided in Supplementary Tables 41 and 42. Quality control was conducted with TPM (transcript per million) matrices visualising read distributions, ridge plots, and scatter plots of sample correlations. Genes with TPM > 2 were defined as expressed in each developmental stage. Gene expression across all samples was normalised using DESeq2 (111), and correlation coefficients were calculated to elucidate relationships among developmental stages. For *O. fusiformis*, batch effects between the two replicates were removed using the limma package (112). To identify differentially expressed genes (DEGs) between adjacent developmental stages, pairwise comparisons were performed using DESeq2 (111), and DEGs were visualised with EnhancedVolcano (113). To recover subtle changes and allow the inclusion of all differences in expression between stages, the log2(fold change) cutoff was set to 0.

### Gene clustering

To elucidate genes with similar expression patterns, we applied soft clustering using the Mfuzz package (114), clustering gene expression profiles based on the average normalised values from DESeq2. To determine the optimal number of clusters, we applied the “centroid” clustering method, “Euclidean” distance with ”ch” index by the R package NbClust (version 3.0.1) (115). Hierarchical clustering was also performed to resolve relationships among developmental stages. Clusters were categorised according to their expression profiles, particularly the stages of highest expression, along with insights from hierarchical clustering and PCA analyses. Gene expression dynamics across the whole profile and within individual clusters were visualised using ComplexHeatmap (116, 117) and line plots. Furthermore, each group was characterised for transcription factor composition based on TF families, and the results were visualised numerically and proportionally using Nightingale rose charts.

### Interspecies RNA-seq comparisons

To evaluate cluster similarity between species (*O. fusiformis*, *C. teleta*, and *C. gigas*), we considered all transcripts within each cluster and orthologous genes identified by OrthoFinder (118) (including one-to-one, one-to-many, and many-to-many relationships), and compute hypergeometric statistics to assess similarity. The results were visualised as a heatmap. We employed Jensen-Shannon Distance (JSD) (119, 120) to estimate inter-sample distances to explore sample relationships across developmental stages between two species. We first identified all one-to-one orthologous genes between the two species and extracted their corresponding TPM values. These TPM values were then quantile transformed with the QuantileTransformer Python script to convert them into probability distributions. We calculated raw JSD values for each pair of developmental stages across species using these distributions. To ensure comparability of JSD values across different stages, we normalised the raw JSD values using a method described in (31). The raw JSD values were visualised in a heatmap, while the normalised values were presented in a line plot alongside the raw data.

### GO and KEGG enrichment

We performed GO term enrichment analyses using the TopGO package (121) to explore each cluster’s possible biological functions. The GO universe was defined based on the annotations in Supplementary Tables 43 and 44. Enrichment statistics were calculated using the Fisher test, and the top 10 or 15 enriched GO terms for each cluster were selected for visualisation. In KEGG pathway enrichment analyses, we first retrieved pathway names using the KO IDs listed in Supplementary Tables 45 and 46 with the KEGGREST package (122). After constructing the KEGG annotation table, we performed enrichment analysis using the ClusterProfiler package (123), setting a *p*-value cutoff of 0.05 and a *q*-value cutoff of 0.5 without *p*-value adjustment. The results of KEGG pathway enrichment were presented in dot plots for clarity and interpretation.

### Phylostratigraphy analysis

To investigate the contribution of evolutionary genes to development, we used available gene ages based on previous gene family evolutionary analyses (31). The proportion of genes within each phylostratum was calculated according to their developmental origins across the entire gene profile and visualised using bar plots. To assess the abundance of these genes across developmental stages, we performed quantile normalisation for each stage, and the normalised values were presented in a heatmap to illustrate their distribution and trends.

### Codon usage

To determine whether there is a bias in codon usage among different clusters, we extracted the transcripts within each cluster and calculated their codon usage individually with the R packages biostrings, seqinR, and coRdon (124–126). To identify changes in codon usage, pairwise comparisons were performed for each codon, with the log_2_(Ratio) used to represent the magnitude of change between clusters. These changes were visualised in bar plots, where, for *O. fusiformis*, codons on the left side of the plot were identified as unstable, while those on the right were classified as stable.

### Gene cloning and *in situ* hybridisation

To generate riboprobes for whole-mount in situ hybridisation and gene expression analyses, the coding sequences of candidate genes were extracted from the genomes of both species, and gene-specific primers were designed via the Primer3 web tool (https://primer3.ut.ee) to amplify gene products ranging from 1000 to 1500 bp. The genes were amplified using cDNA from a mix of developmental stages, and the amplicons were validated by Sanger sequencing. Fixation and whole-mount *in situ* hybridisation (ISH) were performed according to previously published protocols (16, 21, 74, 109).

### Immunostaining

Fixation and antibody staining were conducted as described elsewhere (21). The primary antibodies mouse anti-acetylated α-tubulin (clone 6-11B-1, Merk-Sigma, #MABT868, 1:800) and mouse anti-beta-tubulin (E7, Developmental Studies Hybridoma Bank, 1:20) were diluted in 5% normal goat serum (NGS) in phosphate-buffered saline with 0.5% Triton X-100 (PTx) and incubated overnight at 4 °C. After several washes in 1% bovine serum albumin (BSA) in PTx, samples were incubated with AlexaFluor conjugated secondary antibodies (ThermoFisher Scientific, 1:600) plus DAPI (stock 2 mg/ml, 1:2000) diluted in 5% NGS in PTx overnight at 4 °C.

### Imaging

Representative embryos were cleared and mounted in 70% glycerol in phosphate-buffered saline. Whole-mount in situ hybridisation samples were imaged with a Leica DMRA2 upright microscope equipped with an Infinity5 camera (Lumenera) using differential interference contrast (DIC) optics. Confocal laser scanning microscopy (CLSM) images were taken with a Leica Stellaris 8. CLSM Z-stack projections were built with ImageJ2 (127) and Nikon NIS-elements software. DIC images were digitally stacked with Helicon Focus 7 (HeliconSoft). Brightness and contrast were edited with Adobe Photoshop CC, and figures were built with Adobe Illustrator CC (Adobe Inc.).

## Data availability

The new sequencing data generated in this project have been deposited at the Gene Expression Omnibus portal with BioProject accession number PRJNA1019281.

## Author contributions

YL and JMMD conceived and designed this study. YL and AMCB collected the samples. YL and JMMD performed computational analyses. JW, YL, YK, and AMCB performed gene expression experiments. JW and AMCB conducted immunostaining and imaging analyses. YL and JMMD drafted the text, and all authors read and commented on the manuscript.

## Acknowledgements

We thank members of the Martín-Durán lab for their support and comments on this manuscript. This study employed computational resources from the high-performance computing facility Apocrita at Queen Mary University of London. BBSRC grant BB/W019698/1 supported the research reported in this paper. This work was also funded by the European Union Horizon 2020 Framework Programme (European Research Council Starting Grant agreement number 801669) and the Biotechnology and Biological Sciences Research Council (BB/Y004221/1) to JMMD. JW is supported by a China Scholarship Council doctoral fellowship (CSC NO. 202306330025), and AMCB is supported by a Biotechnology and Biological Sciences Research Council grant (BB/Y004221/1).

