## Supplementary Figures for "Cell fate specification modes shape transcriptome evolution in the highly conserved spiral cleavage"

**Index**

- Supplementary Figures 1 – 14
- Supplementary Tables 1 – 46 (as an Excel spreadsheet)

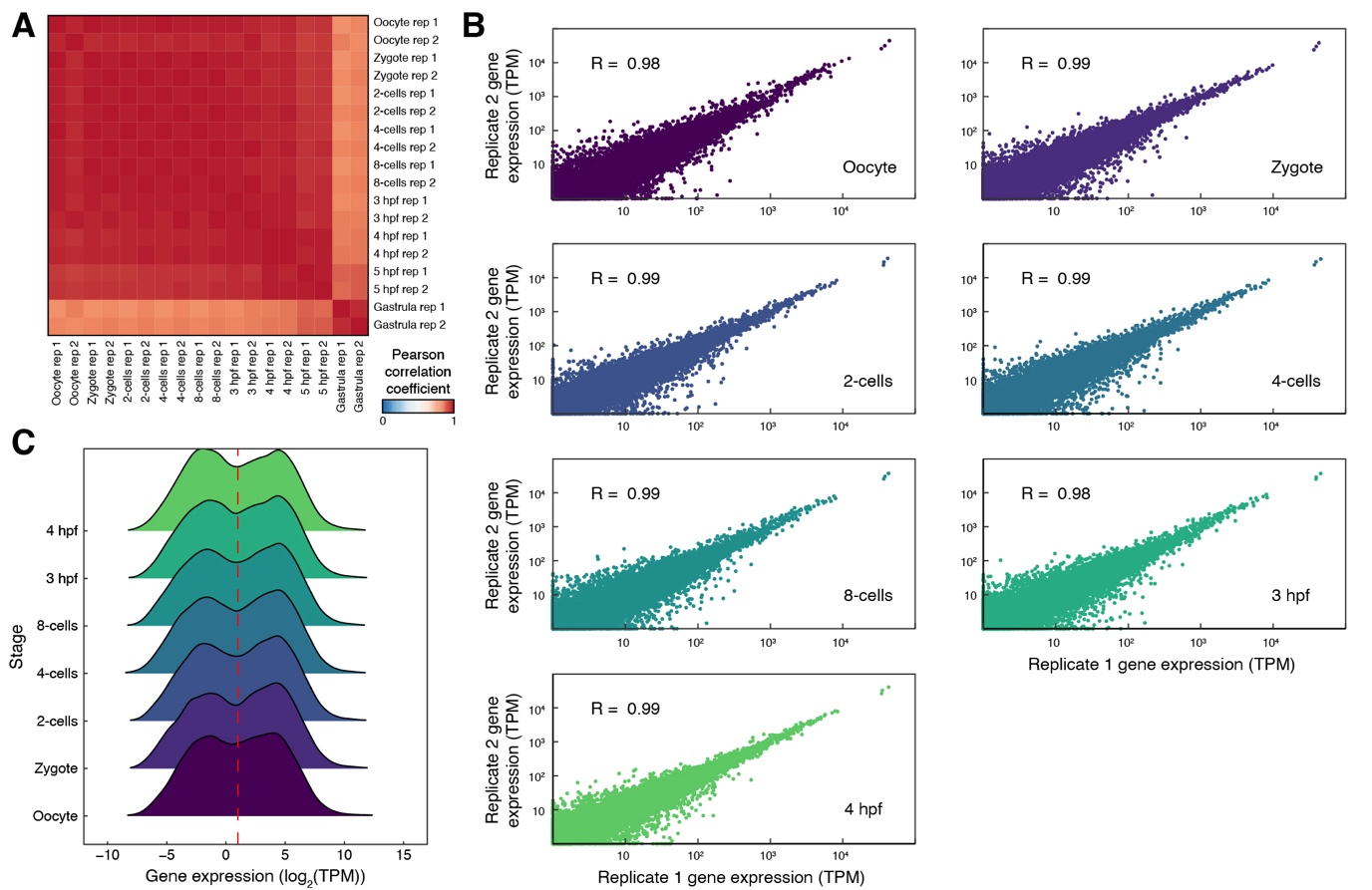

**Supplementary Figure 1 – Quality check of the transcriptomic time course during *O. fusiformis* spiral cleavage.** (**A**) Pearson correlation coefficient matrix of the samples. Cleavage stages are highly similar and different from the gastrula. (**B**) Dot plots of replicate-to-replicate correlation. Replicates for all stages are highly correlated. (**C**) Distribution plots of transcript per million (TPM) values during early spiral cleavage. TPM = 2 (vertical red dotted line) appears to demarcate the transition between expressed and non-expressed.

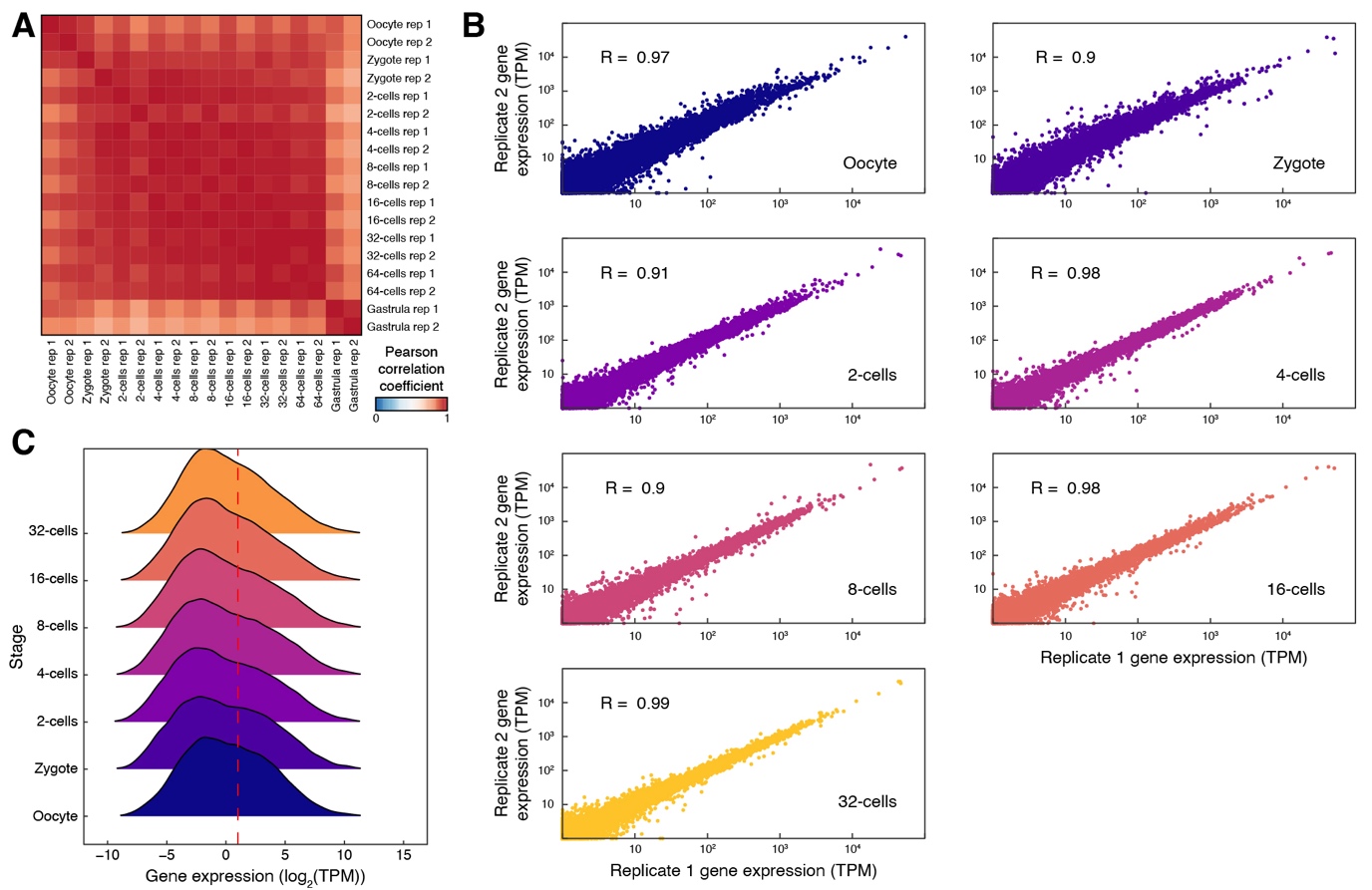

**Supplementary Figure 2 – Quality check of the transcriptomic time course during *C. teleta* spiral cleavage.** (**A**) Pearson correlation coefficient matrix of the samples. Cleavage stages are highly similar and different from the gastrula. (**B**) Dot plots of replicate-to-replicate correlation. Replicates for all stages are highly correlated. (**C**) Distribution plots of transcript per million (TPM) values during early spiral cleavage. TPM = 2 (vertical red dotted line) appears to demarcate the transition between expressed and non-expressed.

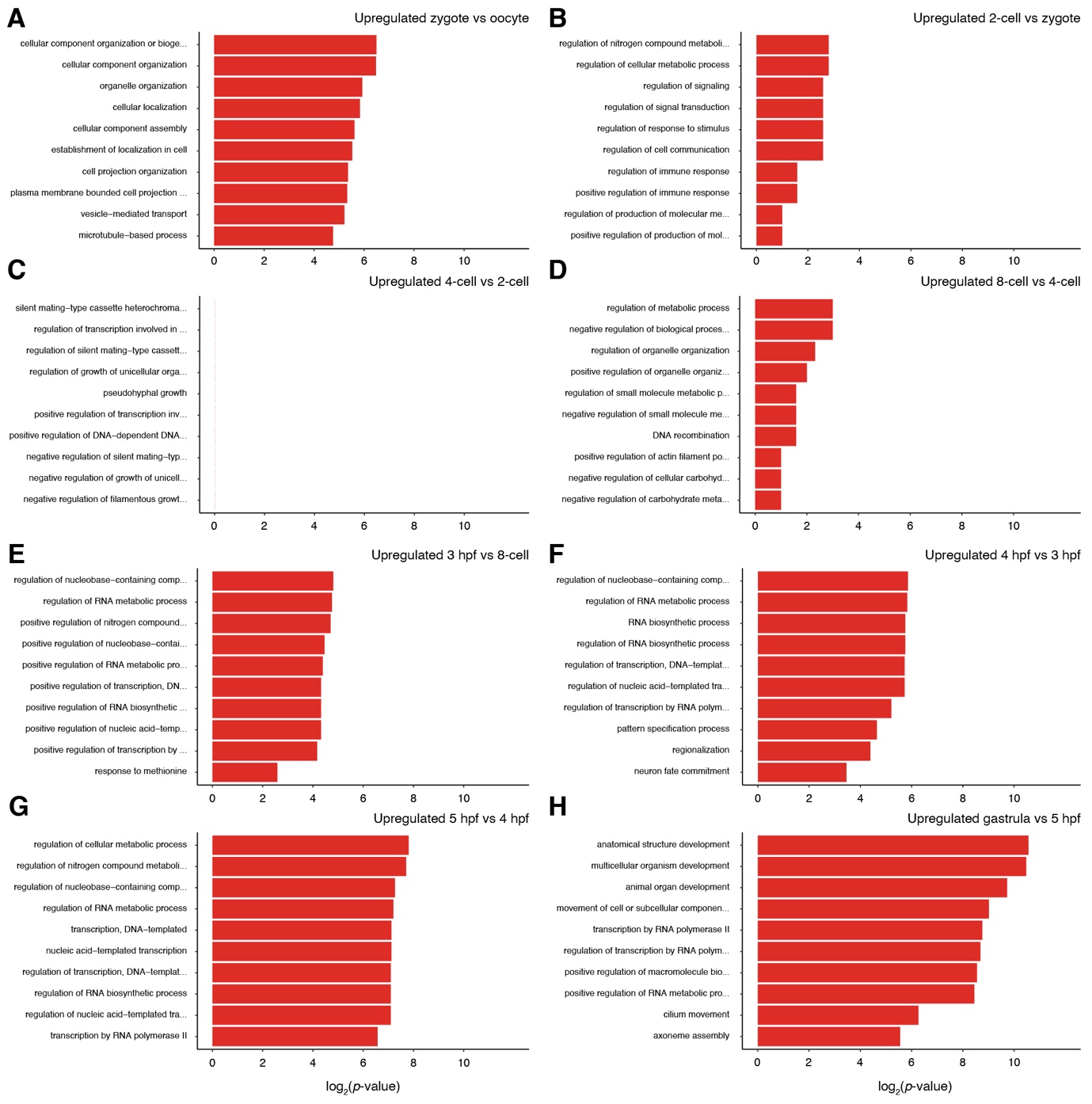

**Supplementary Figure 3 – Gene Ontology enrichment for upregulated genes in *O. fusiformis*.** (**A**–**H**) Bar plots indicating the top ten Gene Ontology (GO) categories amongst upregulated genes in each consecutive pairwise comparison during spiral cleavage in *O. fusiformis*.

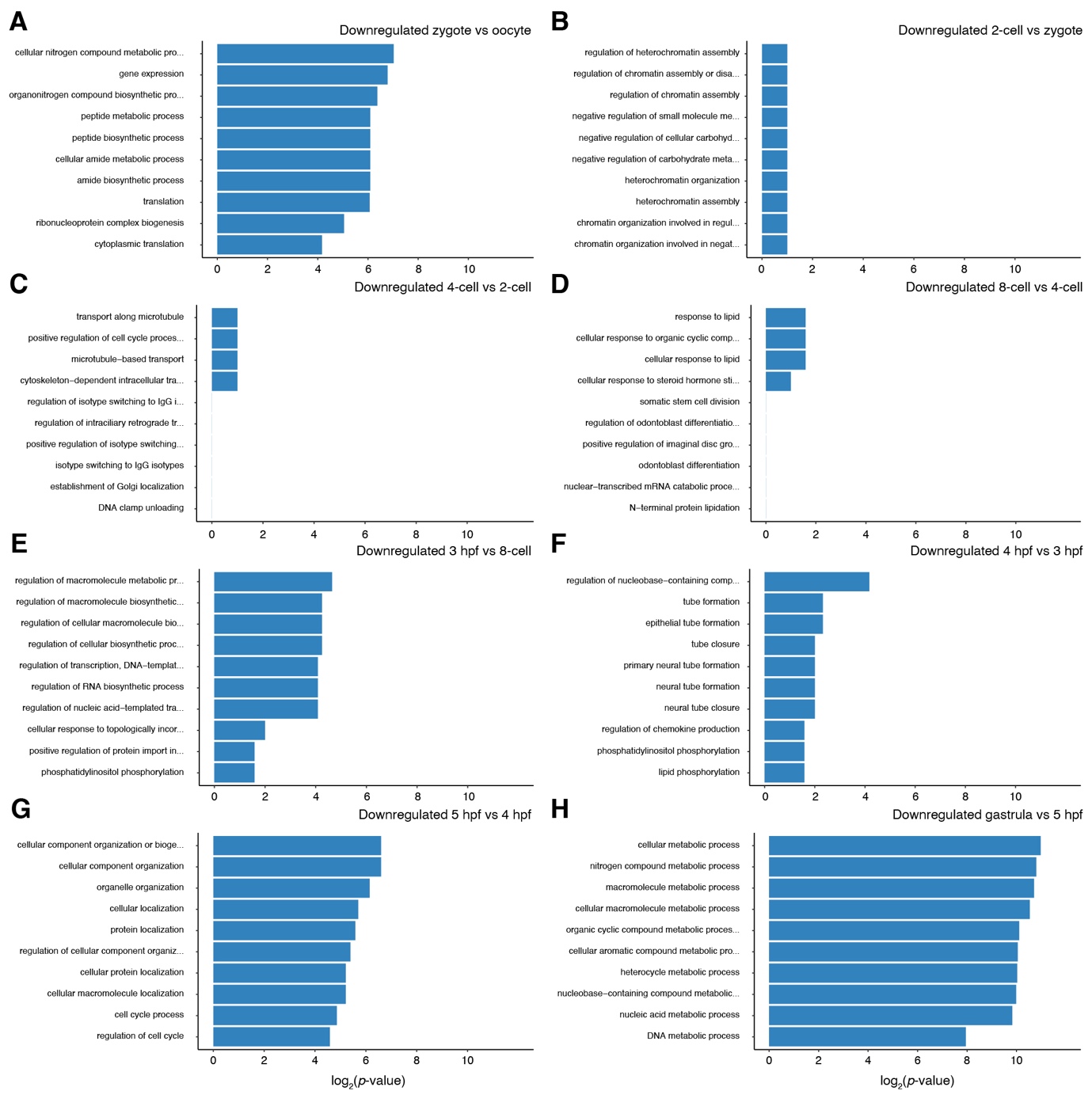

**Supplementary Figure 4 – Gene Ontology enrichment for downregulated genes in *O. fusiformis*.** (**A**–**H**) Bar plots indicating the top ten Gene Ontology (GO) categories amongst downregulated genes in each consecutive pairwise comparison during spiral cleavage in *O. fusiformis*.

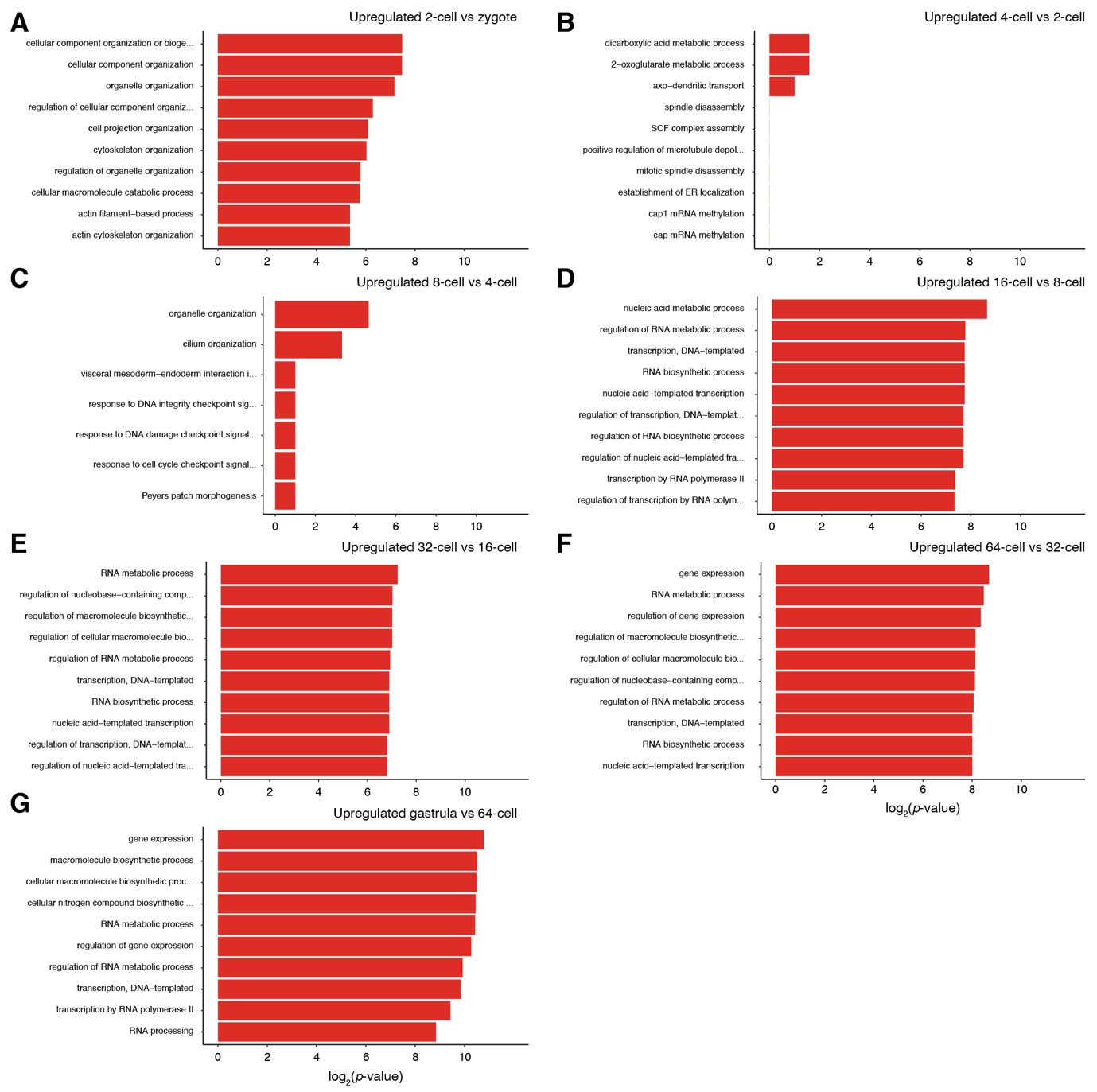

**Supplementary Figure 5 – Gene Ontology enrichment for upregulated genes in *C. teleta*.** (**A**–**G**) Bar plots indicating the top ten Gene Ontology (GO) categories amongst upregulated genes in each consecutive pairwise comparison during spiral cleavage in *C. teleta*.

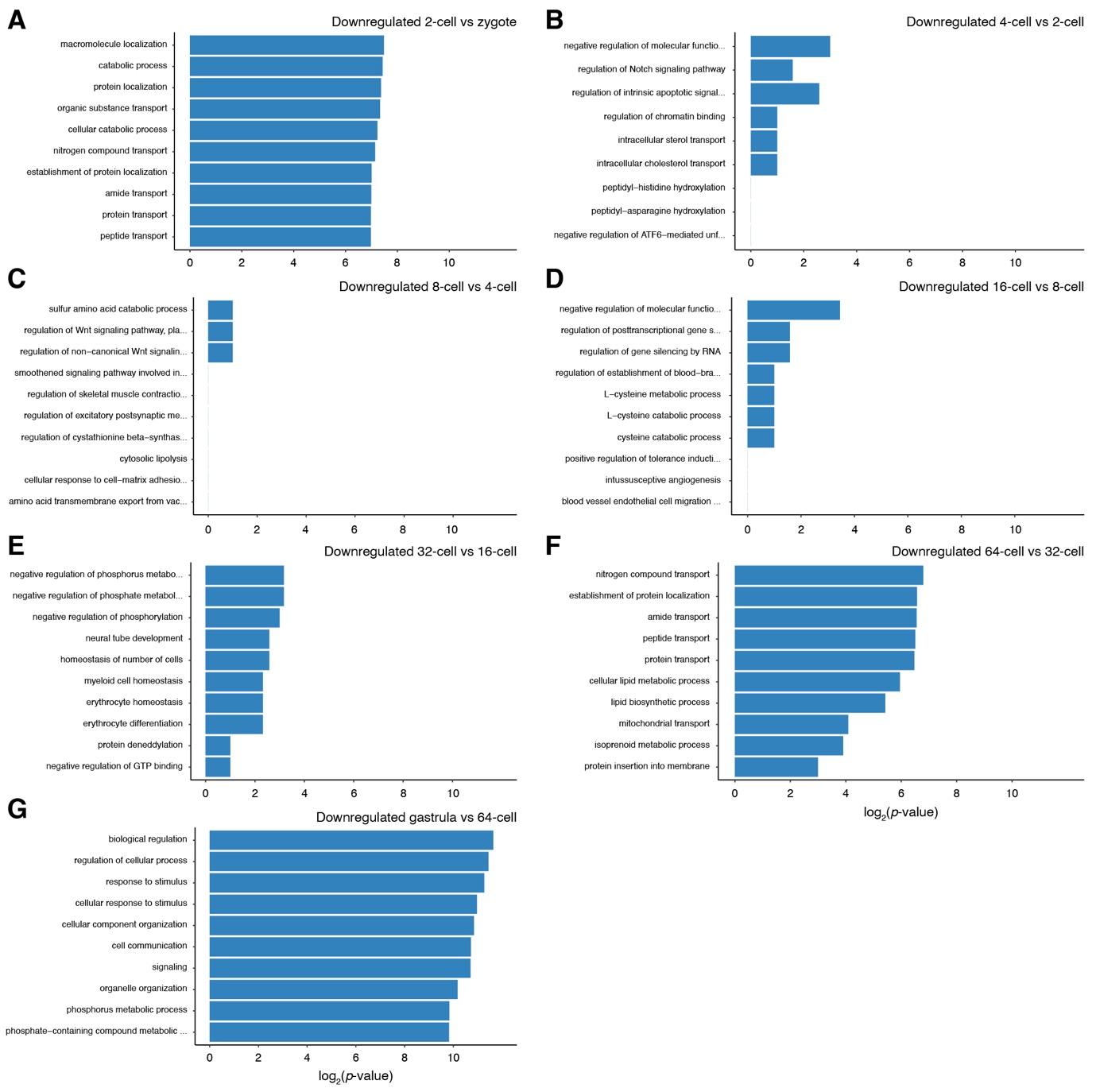

**Supplementary Figure 6 – Gene Ontology enrichment for downregulated genes in *C. teleta*.** (**A**–**G**) Bar plots indicating the top ten Gene Ontology (GO) categories amongst downregulated genes in each consecutive pairwise comparison during spiral cleavage in *C. teleta*.

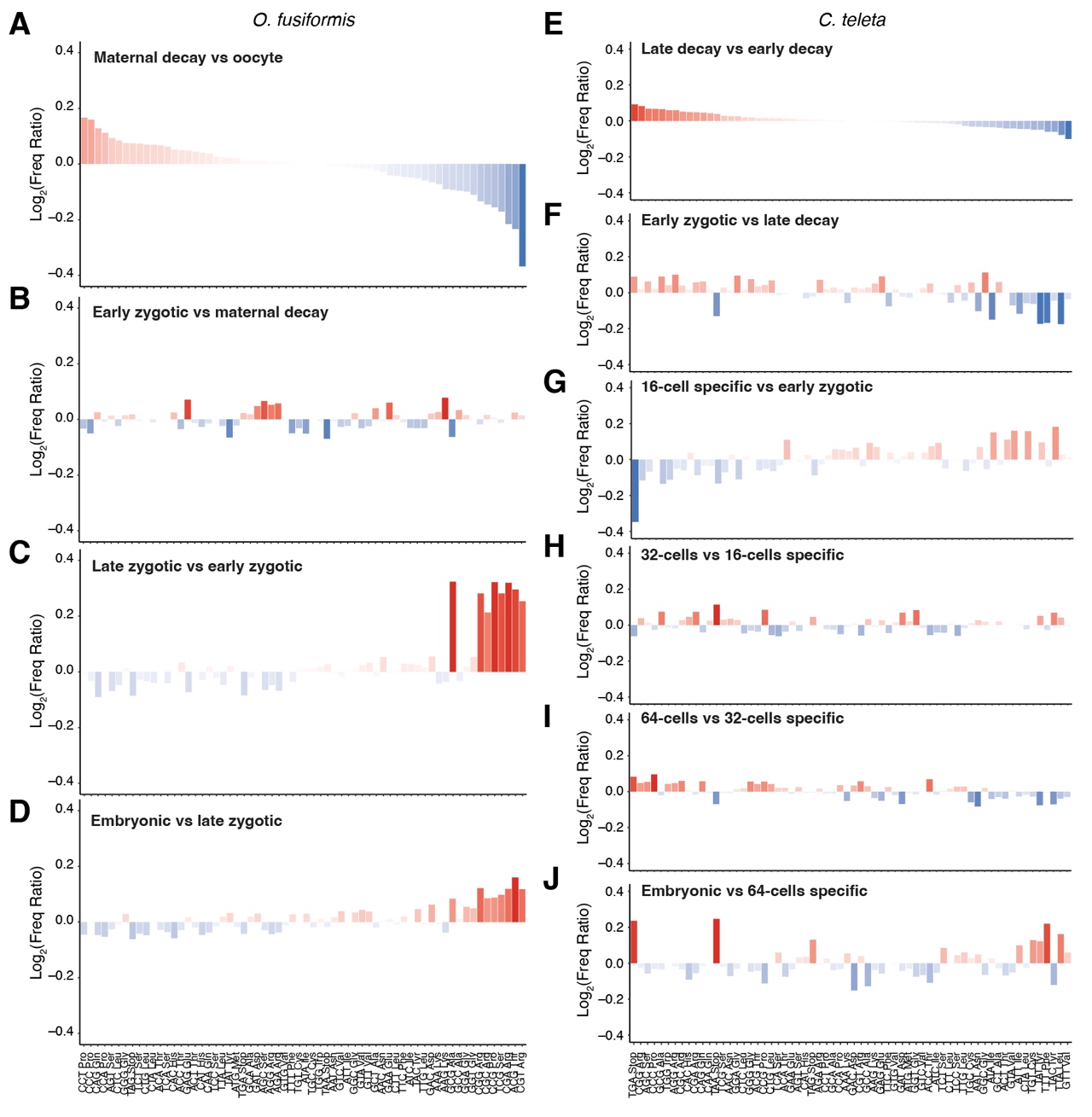

**Supplementary Figure 7 – Codon usage during spiral cleavage.** (**A**–**J**) Bar plots indicating positive (red) or negative (blue) biases in codon usage between pairwise comparisons of clusters of temporal coregulated genes in *O. fusiformis* (**A**–**D**) and *C. teleta* (**E**–**J**). In *O. fusiformis*, codon usage distribution in oocyte-specific genes (**A**) differs from codon usage zygotic genes (**C**, **D**). These marked differences are not that obvious in *C. teleta* (**E**–**J**).

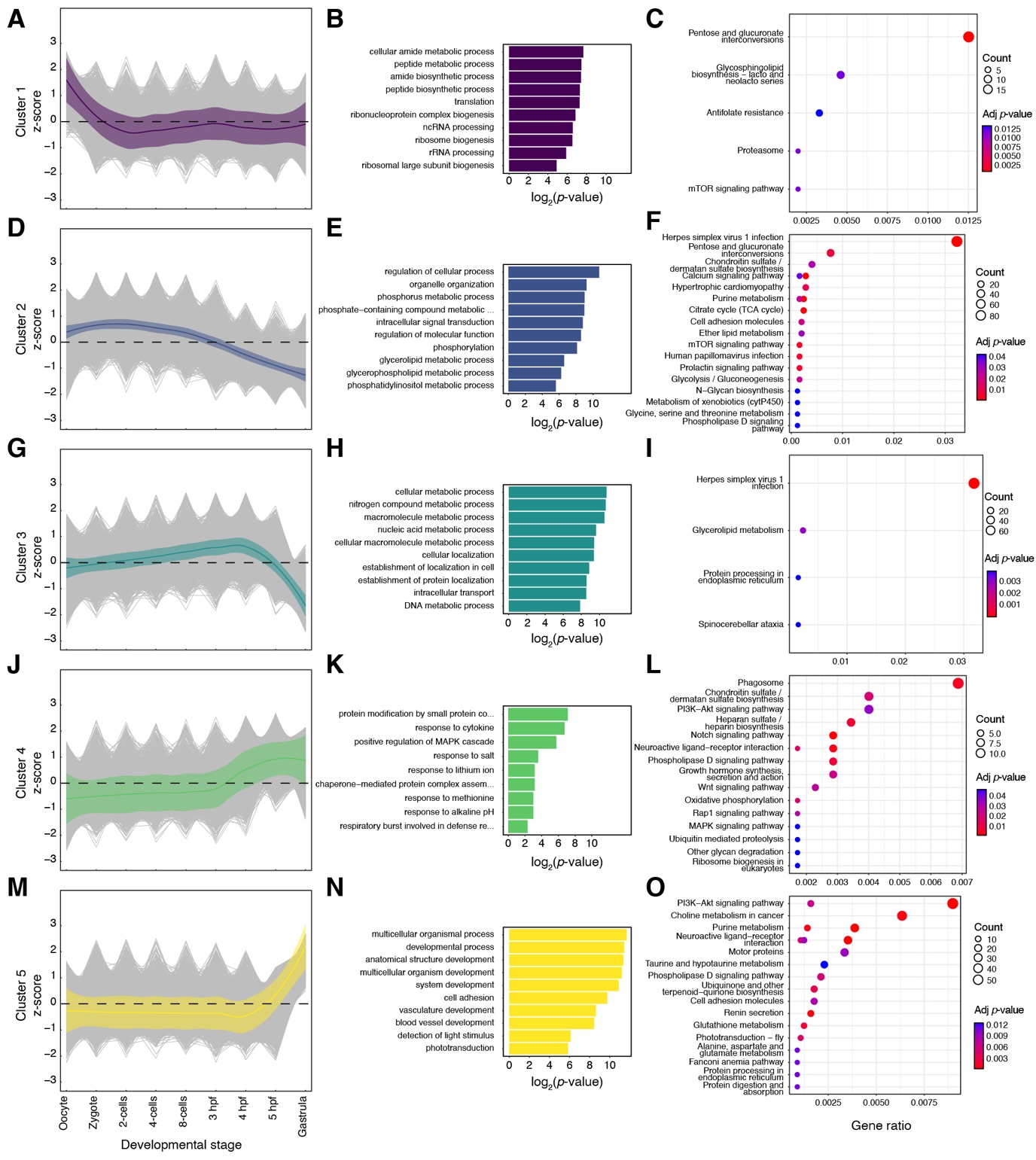

**Supplementary Figure 8 – Functional annotation of clusters of temporally co-regulated genes in *O. fusiformis*.** (**A**, **D**, **G**, **J**, **M**) Gene-wise expression dynamics (grey lines) and locally estimated scatterplot smoothing (coloured lines) for each cluster of temporally coregulated genes during *O. fusiformis* spiral cleavage. Coloured shaded areas represent the standard error of the mean. (**B**, **E**, **H**, **K**, **N**) Bar plots indicating the top ten Gene Ontology (GO) categories amongst genes in each cluster. (**C**, **F**, **I**, **L**, **O**) KEGG enrichment plots for each cluster of coregulated genes during spiral cleavage in *O. fusiformis*.

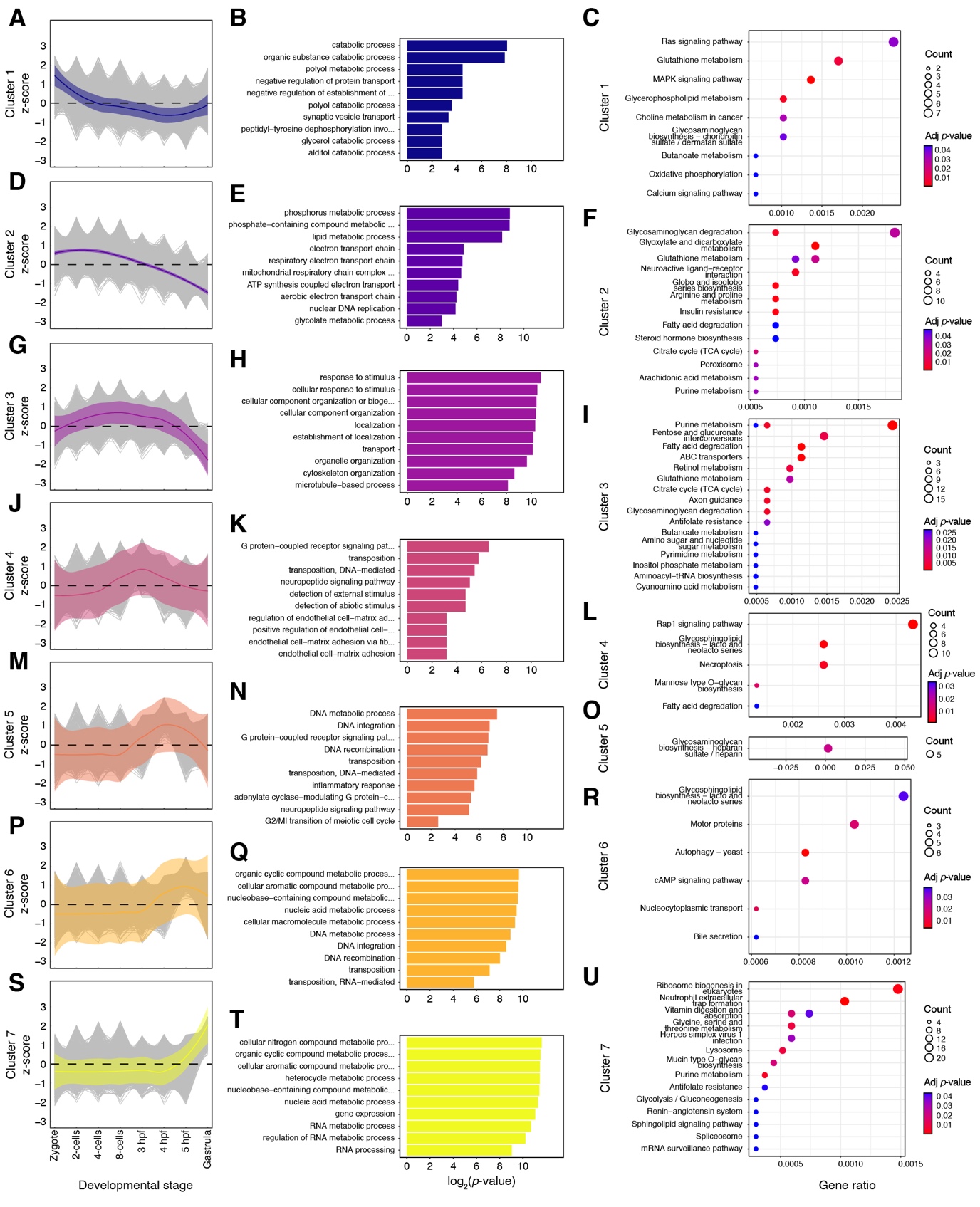

**Supplementary Figure 9 – Functional annotation of clusters of temporally co-regulated genes in *C. teleta*.** (**A**, **D**, **G**, **J**, **M**, **P**, **S**) Gene-wise expression dynamics (grey lines) and locally estimated scatterplot smoothing (coloured lines) for each cluster of temporally coregulated genes during *C. teleta* spiral cleavage. Coloured shaded areas represent the standard error of the mean. (**B**, **E**, **H**, **K**, **N**, **Q**, **T**) Bar plots indicating the top ten Gene Ontology (GO) categories amongst genes in each cluster. (**C**, **F**, **I**, **L**, **O**, **R**, **U**) KEGG enrichment plots for each cluster of coregulated genes during spiral cleavage in *C. teleta*.

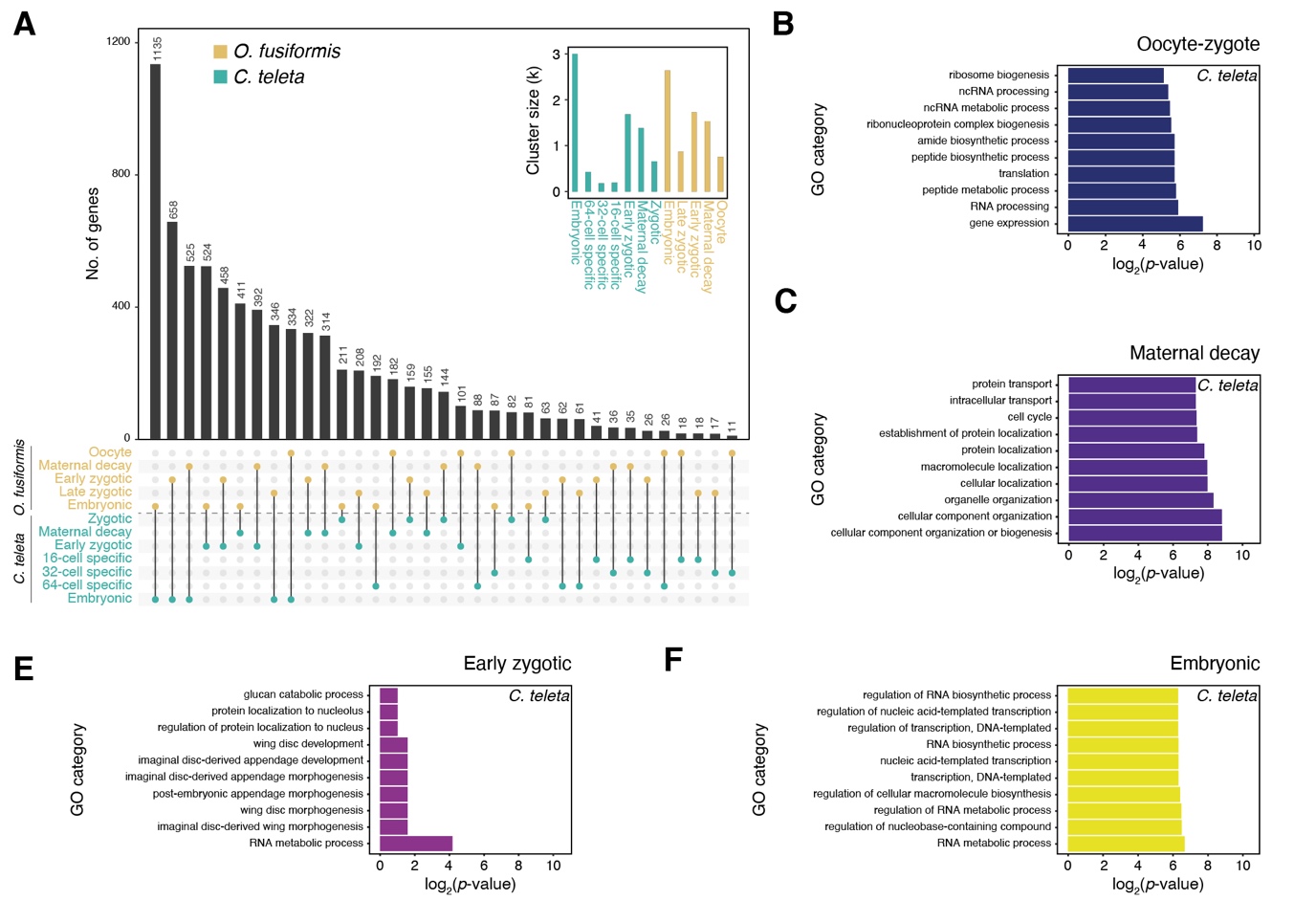

**Supplementary Figure 10 – The comparison of clusters of temporally co-regulated genes between conditional and autonomous spiral cleavage.** (**A**) Upset plot indicating the number of shared one-to-one orthologs between clusters of temporally coregulated genes. The inset indicates the total number of genes in each cluster. (**B**–**F**) Bar plots indicating the top ten Gene Ontology (GO) categories amongst shared orthologous genes in the oocyte/zygote, maternal decay, early zygotic, and embryonic clusters, according to the GO annotation for the *C. teleta* ortholog.

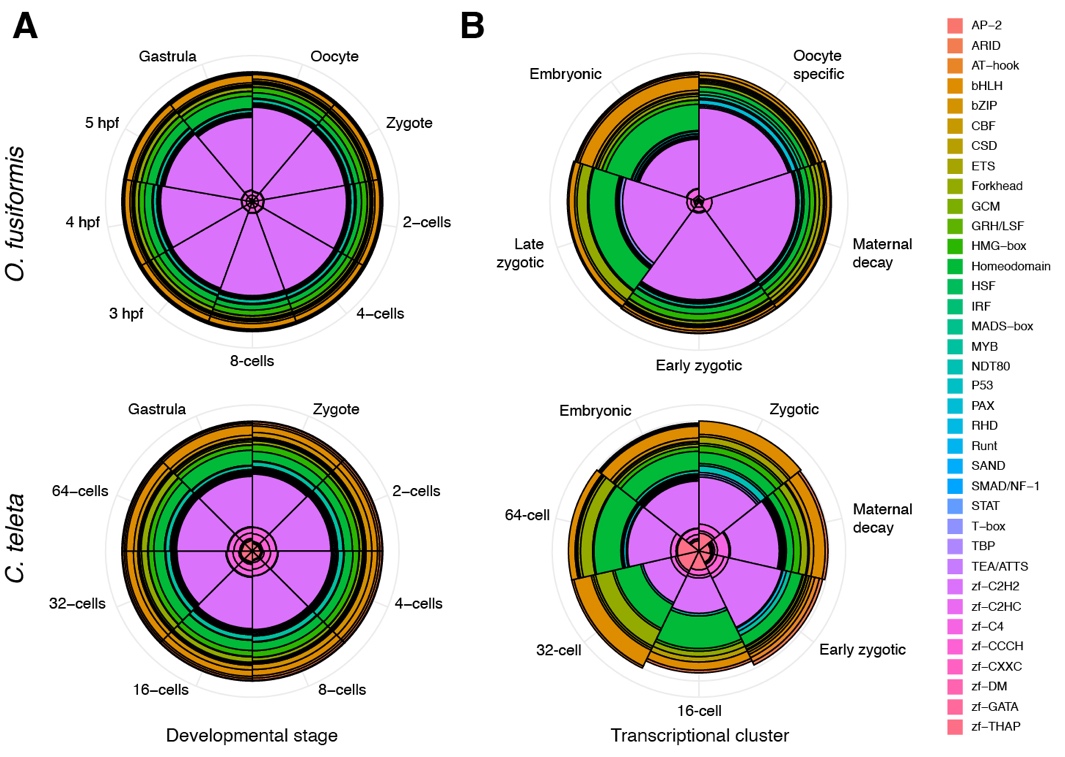

**Supplementary Figure 11 – The proportion of transcription factors during spiral cleavage.** (**A**, **B**) Rose plots of transcription factor (TF) distribution (as a percentage) according to developmental time point (**A**) and cluster of coregulated genes (**B**).

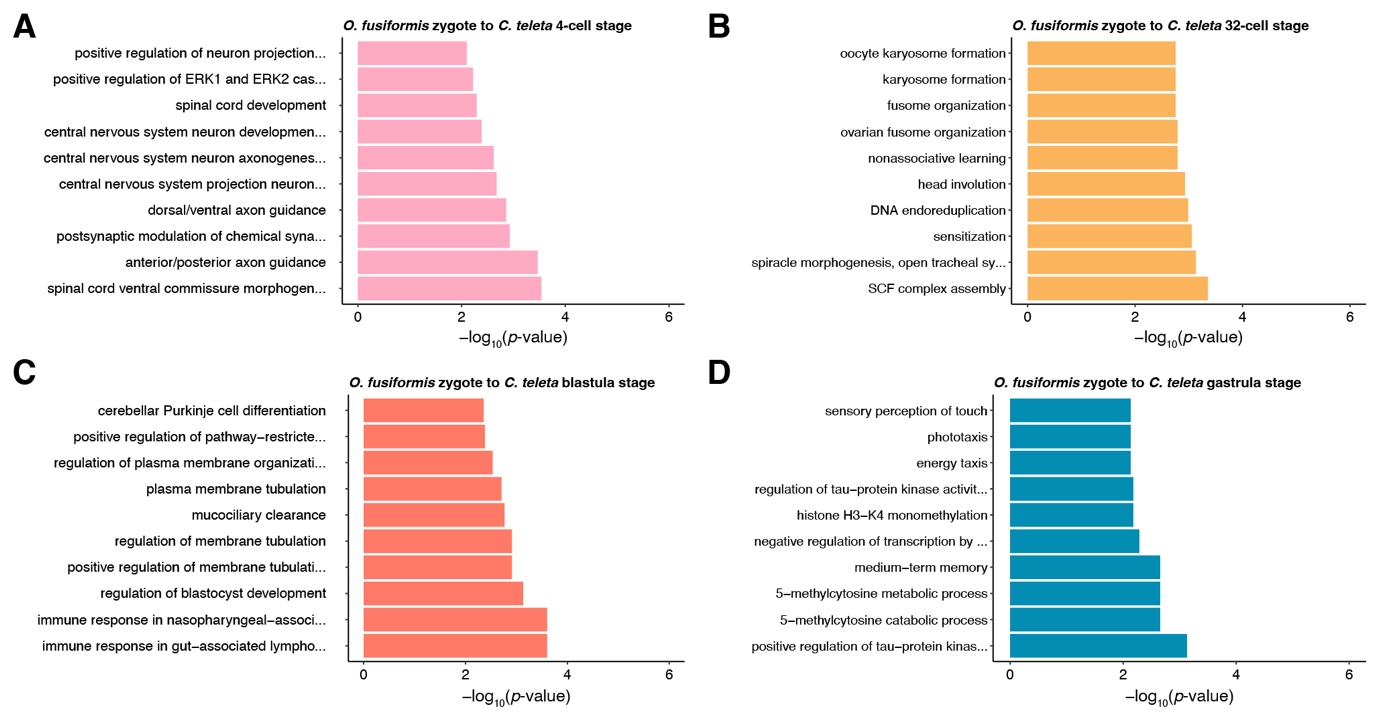

**Supplementary Figure 12 – Gene Ontology enrichment in maternal genes in *O. fusiformis*.** (**A**–**D**) The bar plots indicate the top ten Gene Ontology (GO) categories amongst maternal genes in *O. fusiformis* that are expressed later in *C. teleta*.

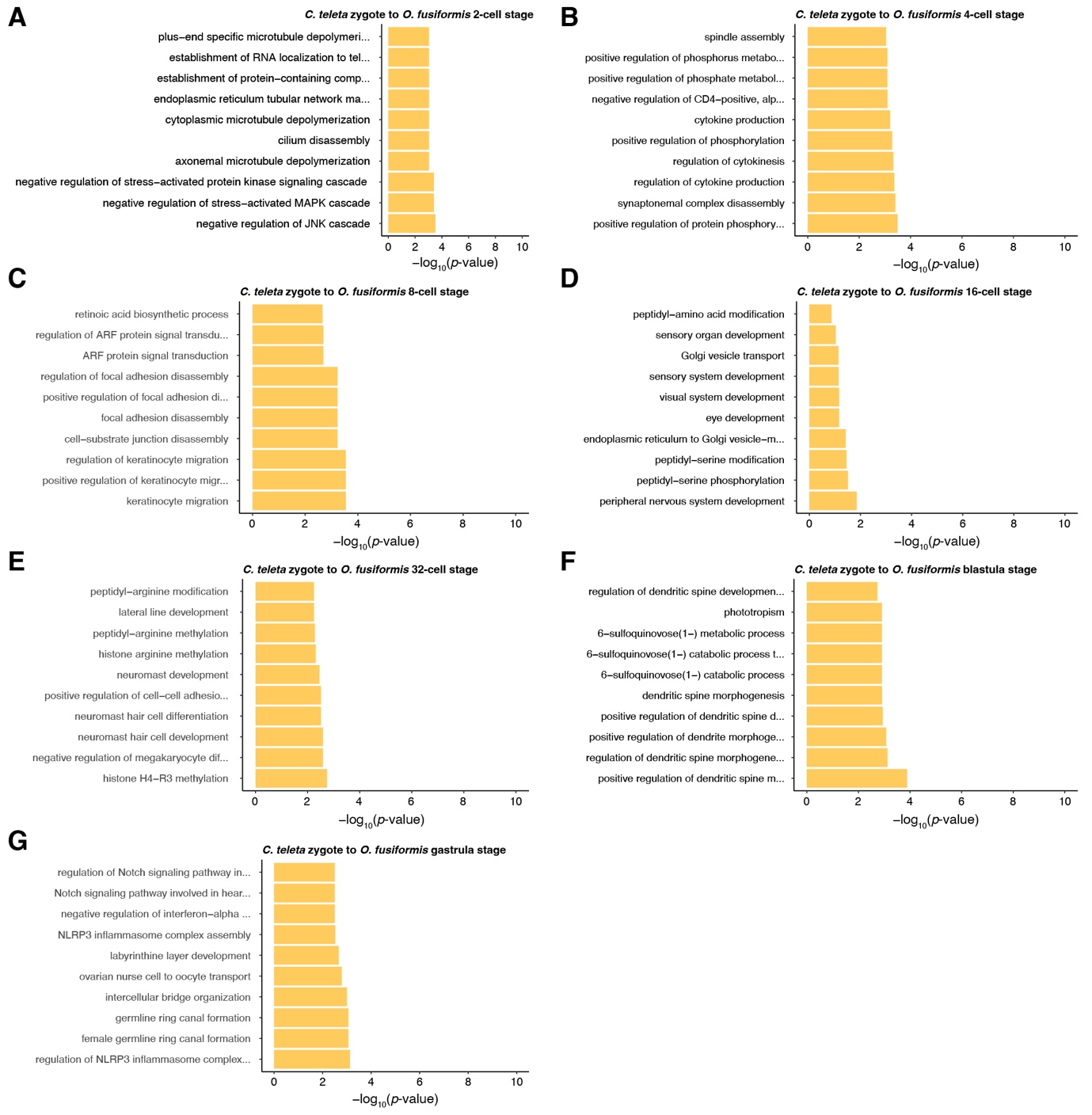

**Supplementary Figure 13 – Gene Ontology enrichment in maternal genes in *C. teleta*.** (**A**–**G**) The bar plots indicate the top ten Gene Ontology (GO) categories amongst maternal genes in *C. teleta* that are expressed later in *O. fusiformis*.

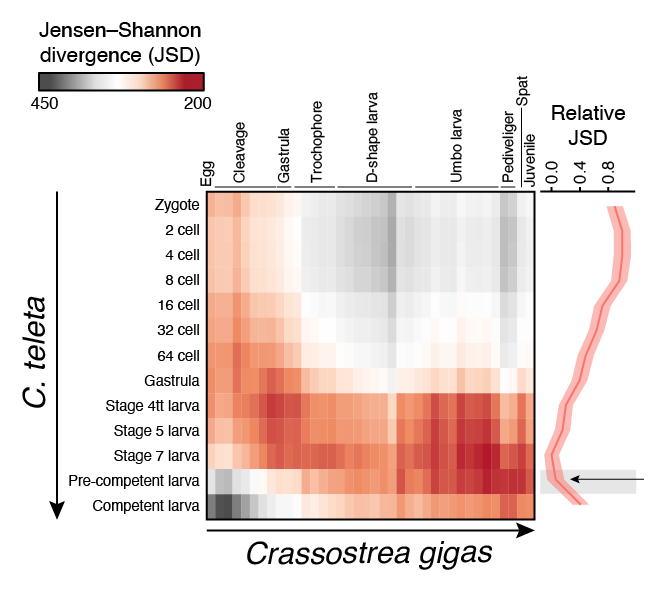

**Supplementary Figure 14 – Transcriptional similarity between the development of the annelid *Capitella teleta* and the bivalve *Crassostrea gigas*.** Jensen-Shannon transcriptomic divergence between all possible inter-species pairwise comparisons during the entire life cycle, from oocyte to juvenile or competent larva of *C. teleta* and *Crassostrea gigas*.

**List of Supplementary Tables**

**Supplementary Table 1 –** Number of up- and downregulated genes during early embryogenesis in *O. fusiformis* and *C. teleta*.

**Supplementary Table 2 –** Upregulated genes between the zygote and oocyte in *O. fusiformis*.

**Supplementary Table 3 –** Downregulated genes between the zygote and oocyte in *O. fusiformis*.

**Supplementary Table 4 –** Upregulated genes between the 2-cell stage and the zygote in *O. fusiformis*.

**Supplementary Table 5 –** Downregulated genes between the 2-cell stage and the zygote in *O. fusiformis*.

**Supplementary Table 6 –** Upregulated genes between the 4- and the 2-cell stages in *O. fusiformis*.

**Supplementary Table 7 –** Downregulated genes between the 4- and the 2-cell stages in *O. fusiformis*.

**Supplementary Table 8 –** Upregulated genes between the 8- and the 4-cell stages in *O. fusiformis*.

**Supplementary Table 9 –** Downregulated genes between the 8- and the 4-cell stages in *O. fusiformis*.

**Supplementary Table 10 –** Upregulated genes between 3 hpf and the 8-cell stages in *O. fusiformis*.

**Supplementary Table 11 –** Downregulated genes between 3 hpf and the 8-cell stages in *O. fusiformis*.

**Supplementary Table 12 –** Upregulated genes between 4 and 3 hpf in *O. fusiformis*.

**Supplementary Table 13 –** Downregulated genes between 4 and 3 hpf in *O. fusiformis*.

**Supplementary Table 14 –** Upregulated genes between 5 and 4 hpf in *O. fusiformis*.

**Supplementary Table 15 –** Downregulated genes between 5 and 4 hpf in *O. fusiformis*.

**Supplementary Table 16 –** Upregulated genes between the gastrula stage and 5 hpf in *O. fusiformis*.

**Supplementary Table 17 –** Downregulated genes between the gastrula stage and 5 hpf in *O. fusiformis*.

**Supplementary Table 18 –** Upregulated genes between the 2-cell stage and the zygote in *C. teleta*.

**Supplementary Table 19 –** Downregulated genes between the 2-cell stage and the zygote in *C. teleta*.

**Supplementary Table 20 –** Upregulated genes between the 4- and the 2-cell stages in *C. teleta*.

**Supplementary Table 21 –** Downregulated genes between the 4- and the 2-cell stages in *C. teleta*.

**Supplementary Table 22 –** Upregulated genes between the 8- and the 4-cell stages in *C. teleta*.

**Supplementary Table 23 –** Downregulated genes between the 8- and the 4-cell stages in *C. teleta*.

**Supplementary Table 24 –** Upregulated genes between the 16- and the 8-cell stages in *C. teleta*.

**Supplementary Table 25 –** Downregulated genes between the 16- and the 8-cell stages in *C. teleta*.

**Supplementary Table 26 –** Upregulated genes between the 32- and the 16-cell stages in *C. teleta*.

**Supplementary Table 27 –** Downregulated genes between the 32- and the 16-cell stages in *C. teleta*.

**Supplementary Table 28 –** Upregulated genes between the 64- and the 32-cell stages in *C. teleta*.

**Supplementary Table 29 –** Downregulated genes between the 64- and the 32-cell stages in *C. teleta*.

**Supplementary Table 30 –** Upregulated genes between the gastrula and the 64-cell stage in *C. teleta*.

**Supplementary Table 31 –** Downregulated genes between the gastrula and the 64-cell stage in *C. teleta*.

**Supplementary Table 32 –** Cluster allocation for the expressed transcripts during *O. fusiformis* early embryogenesis.

**Supplementary Table 33 –** Cluster allocation for the expressed transcripts during *C. teleta* early embryogenesis.

**Supplementary Table 34 –** Transcription factor composition by developmental time point in *O. fusiformis*.

**Supplementary Table 35 –** Transcription factor composition by coregulated gene clusters in *O. fusiformis*.

**Supplementary Table 36 –** Transcription factor composition by developmental time point in *C. teleta*.

**Supplementary Table 37 –** Transcription factor composition by coregulated gene clusters in *C. teleta*.

**Supplementary Table 38 –** One-to-one orthologs exhibiting temporal shifts in gene activation during cleavage between *O. fusiformis* and *C. teleta*.

**Supplementary Table 39 –** Comparison in the number of genes exhibiting temporal shifts in gene activation during cleavage between *O. fusiformis* and *C. teleta*.

**Supplementary Table 40 –** Expression values (TPM) of the validated candidate genes during spiral cleavage in *O. fusiformis* and *C. teleta*.

**Supplementary Table 41 –** Sequencing and mapping statistics for *O. fusiformis*.

**Supplementary Table 42 –** Sequencing and mapping statistics for *C. teleta*.

**Supplementary Table 43 –** Gene Ontology universe for *O. fusiformis*.

**Supplementary Table 44 –** Gene Ontology universe for *C. teleta*.

**Supplementary Table 45 –** KEGG universe for *O. fusiformis*.

**Supplementary Table 46 –** KEGG universe for *C. teleta*.
